# Adenine nucleotide translocase 2 (ANT2) deficiency reprograms ferroptosis in alveolar progenitor cells to promote emphysema

**DOI:** 10.64898/2026.07.11.737954

**Authors:** Ugonna Mbaekwe, Jian Shi, Nai-Chun Ting, Qianjiang Hu, Sebastien Gingras, Melanie Königshoff, Corrine R. Kliment

**Affiliations:** Division of Pulmonary, Allergy and Critical Care Medicine, University of Pittsburgh School of Medicine; Pittsburgh, PA 15213; Department of Cellular and Molecular Pathology, University of Pittsburgh School of Medicine; Department of Immunology and Innovative Technologies Development (ITD) Core, University of Pittsburgh School of Medicine; Center for Lung Aging and Regeneration, University of Pittsburgh; Geriatric Research Education and Clinical Center (GRECC) at the VA Pittsburgh Healthcare System, Pittsburgh, Pennsylvania

**Keywords:** Chronic obstructive pulmonary disease (COPD), Adenine Nucleotide Translocase (ANT), Cigarette smoke, Mitochondria, Ferroptosis

## Abstract

Stem cell dysfunction and loss of renewal capacity are primary characteristics of tissue aging and decremental regeneration in response to injury. Alveolar type 2 cells (AT2) are key progenitor cells responsible for lung repair and are thought to be dysfunctional in diseases such as chronic obstructive pulmonary disease (COPD). AT2 cells are highly metabolic and rely on mitochondria, but how mitochondrial mechanisms influence their maintenance and cell fate is unclear. This gap is critical as no current therapies target lung repair or mitochondrial function in COPD. Here, we report that adenine nucleotide translocase 2 (ANT2), a key ATP/ADP transporter, is reduced in AT2 cells from COPD lungs, and that ANT2 loss impairs bioenergetics (ATP). We also identify, for the first time, ferroptotic susceptibility as a consequence of ANT2 loss in AT2 cells, leading to impaired self-renewal and progenitor capacity in alveolar organoids. Together, loss of ANT2 and the associated cellular dysfunction resulted in worsened lung damage or emphysema due to cigarette smoke in mice. Therapeutic restoration of ANT2 expression resulted in renewed AT2 stem cell function and prevention of emphysema by reducing oxidative stress and ferroptosis. These findings highlight the importance of ANT2 in metabolic regulation, plasticity, and cell resiliency of AT2 cells in the lung and that ANT2 is a potential target for lung repair.

**Graphical Abstract:** 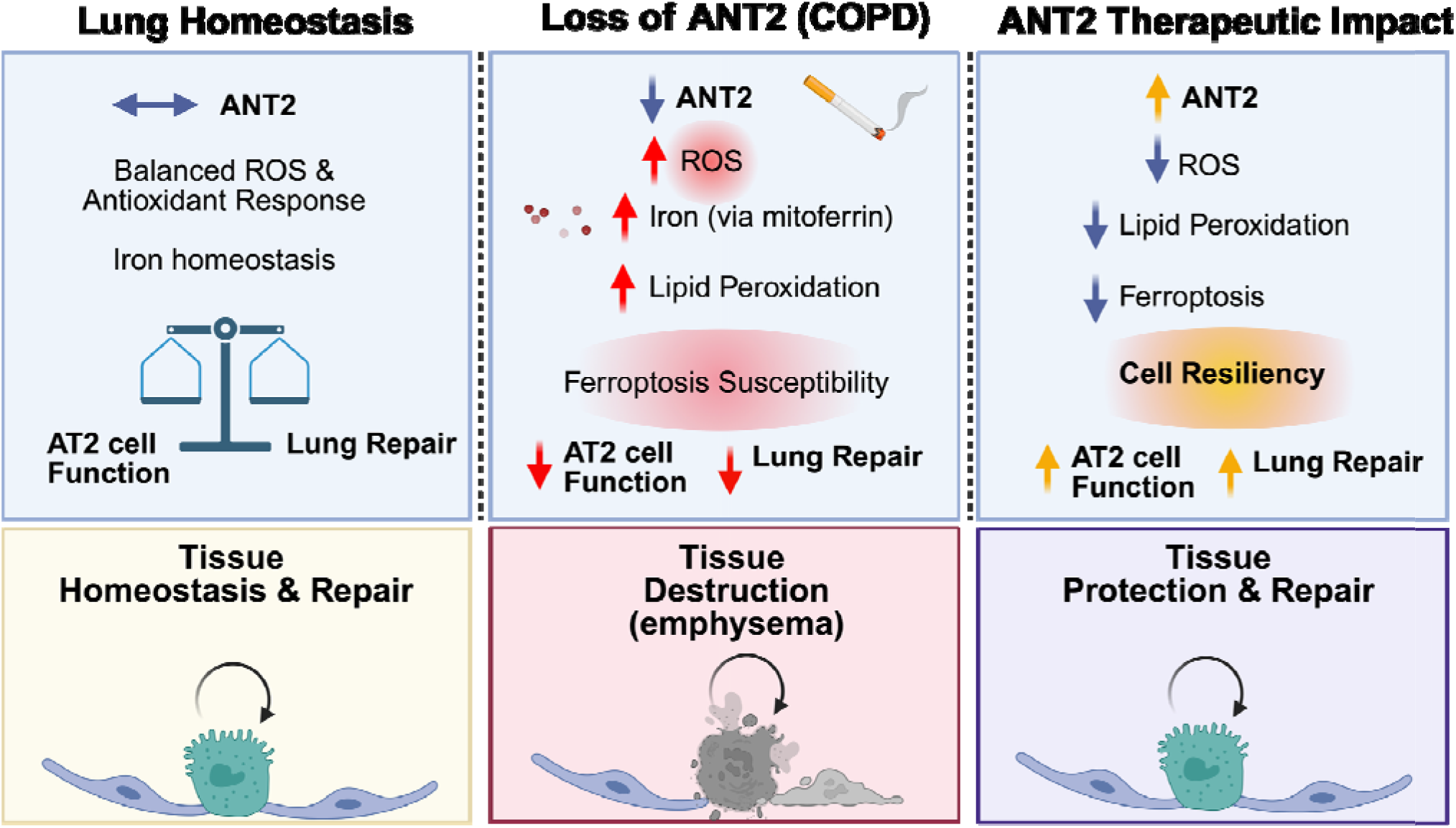

## Introduction

Stem cell dysfunction and loss of renewal capacity are primary characteristics of tissue aging and decremental regeneration in response to injury. Alveolar type 2 cells are a key progenitor cell in the lung and are dysfunctional in chronic lung diseases, such as chronic obstructive pulmonary disease (COPD), which may represent states of accelerated lung aging. Affecting nearly 10.6% of the world population, COPD is the 3^rd^ leading cause of death with no current therapies that reverse disease pathogenesis (1). Risk factors for disease development include cigarette smoke (CS), environmental exposures and genetic alterations (2,3) that contribute to disease pathogenesis through epithelial cell injury, increased oxidative stress and mitochondrial dysfunction (4–8). The alveolar epithelium is composed of several key cell types including alveolar type 1 (AT1) and type 2 (AT2) cells. At homeostasis, AT2 cells will self-renew, and upon injury to the lung, such as from CS, AT2 cells must undergo rapid proliferation and differentiation to AT1 cells to replace damaged lung epithelium (9–12). To accomplish these homeostatic functions, AT2 cells have a high energy demand and rate of mitochondrial oxidative respiration (6,13,14), given their roles in alveolar repair and production of pulmonary surfactant, both essential for lung health.

AT2 cells must maintain plasticity to shift between a state of quiescence to proliferation or differentiation. In a number of stem cell types, mitochondrial metabolism is a strong determinant of this cell behavior and cell fate (15). Cellular proliferation, as occurs with AT2 stem cell self-renewal, has a high energetic demand (ATP and necessary cofactors such as NADPH) (16,17) and relies heavily on the generation of ATP via mitochondrial oxidative phosphorylation and glycolysis (18). Mitochondrial reactive oxygen species (ROS) can force stem cells to proliferate and induce differentiation (19–22), where cells shift metabolism to oxidative phosphorylation. The specific mitochondrial factors that regulate AT2 stem cell function and fate remain unclear.

COPD is defined by the Global Initiative for Chronic Obstructive Lung Disease (GOLD) guidelines as airflow limitation that is not reversible (2). Cigarette smoke (CS) is an important risk factor for COPD development and is known to induce oxidative stress and epithelial cell injury, resulting in increased airway and alveolar inflammation (23–27). The resulting chronic inflammation precipitates further protease destruction of the lung. The production of reactive oxygen species (ROS) has also been suggested as a key mediator in the early epithelial responses and cellular dysfunction due to CS (23–26). Mitochondria are a key source of ROS, raising the prospect of mitochondrial dysfunction as an important driver of COPD (28,29). AT2 cells from human COPD lungs have mitochondrial DNA damage and altered mitochondrial structure and metabolism (5,7,30).

A significant knowledge gap exists regarding how mitochondrial dysfunction promotes ferroptosis in AT2 cells in COPD. Ferroptosis, an iron-dependent form of cell death increasingly associated with COPD, depends on iron availability and the presence of ROS, which drive lipid peroxidation and trigger ferroptotic cell death. *In vitro* studies show that CS exposure drives mitochondrial dysfunction and oxidative stress, thereby increasing labile iron and lipid peroxidation (31,32) and inducing ferroptosis, a non-apoptotic form of cell death, in lung epithelial cells (33). These findings extend to clinical disease: recent research links iron dysregulation and ferroptosis to COPD pathogenesis, initiated by oxidative stress, metabolic alterations, and mitochondrial dysfunction (33–35). Loss of epithelial cells through ferroptosis impairs the lung’s capacity to regenerate and repair tissue, exacerbating disease progression. Ferroptosis inhibitors have demonstrated protective effects against cigarette smoke (CS)-induced ferroptosis in lung epithelial cells (33). While mitochondrial dysfunction has been associated with ferroptosis, the specific mitochondrial drivers of ferroptosis susceptibility in COPD remain unclear. Protecting mitochondrial function may help reduce ferroptosis and enhance cell resilience in chronic lung disease, such as COPD.

Adenine nucleotide translocases (ANT1-4) are abundant ADP/ATP transporters on the inner mitochondrial membrane that are critical for the movement of ATP produced by the electron transport chain (ETC) to the cytoplasm (36,37). ANT2 is highly expressed in rapidly proliferating muscle stem cells and plays a role in their differentiation to myoblasts (38). We have previously demonstrated that ANT1 and ANT2 are reduced in lung tissue from patients with COPD, that loss of ANT2 results in reduced mitochondrial respiration, and that expression of ANT2 is protective in the airway epithelium (39).

In this study, we show that ANT2 is lost in AT2 lung progenitor cells in COPD patients, resulting in reduced progenitor capacity through increased susceptibility to ferroptosis. Loss of ANT2 specifically enhances mitochondrial ROS and iron influx through mitoferrin-1, creating the perfect storm to drive ferroptosis. Lastly, we demonstrate that enhancing ANT2 expression recovers AT2 cell plasticity and reparative function, ultimately protecting the lung from emphysema development. These findings highlight the importance of ANT2 in the metabolic regulation, oxidative state and cell fate of AT2 cells to promote alveolar lung repair.

## Results

### ANT2 is reduced in AT2 cells in human COPD

We have previously shown that gene expression of *SLC25A4* (ANT1) and *SLC25A5* (ANT2) is reduced in the lung tissue of patients with COPD (at all GOLD severity stages) compared to non-smoker control subjects without lung disease (39). We confirmed that ANT1 and ANT2 are reduced at the protein level in whole-lung lysates from subjects with advanced COPD (GOLD Stage 3-4) compared with non-smoker control subjects (**Figure 1A**). Using publicly available single-cell RNA sequencing (scRNA-seq) data of human lung epithelial cells (EpCAM+) isolated from COPD and control subjects, *SLC25A4* (ANT1) and *SLC25A5* (ANT2) are notably enriched in AT2 cells (specifically identified using *SFTPC* and *SFTPB* expression) (**Figure 1B**) and reduced in subjects with COPD (**Figure 1B-F**). ANT1 and ANT2 are also enriched in ionocyte populations and reduced in COPD. Pulmonary ionocytes are rare, comprising only about 0.5-1.5% of the airway epithelial cells, and they exhibit disproportionately high expression of Cystic Fibrosis Transmembrane Conductance Regulator (CFTR) (40,41). We confirmed the reduction of *SLC25A5* expression in AT2 cells in a second publicly available human COPD scRNA-seq dataset from the COPD Cell Atlas (42), which profiled whole lung tissue from patients with and without COPD (**Figure 1G**). Within the lung epithelial cell populations, we observe a similar enrichment of *SLC25A5* (ANT2) in AT2 cells and a significant reduction in AT2 cells of subjects with COPD. We next stained human lung tissue and colocalized ANT1 and ANT2 protein expression in AT2 cells (HTII-280^+^) in human lung tissue from control and COPD subjects, demonstrating that both ANT1 and ANT2 are reduced in AT2 cells at the protein level (**Figure 1H, I**).

**Figure 1:**
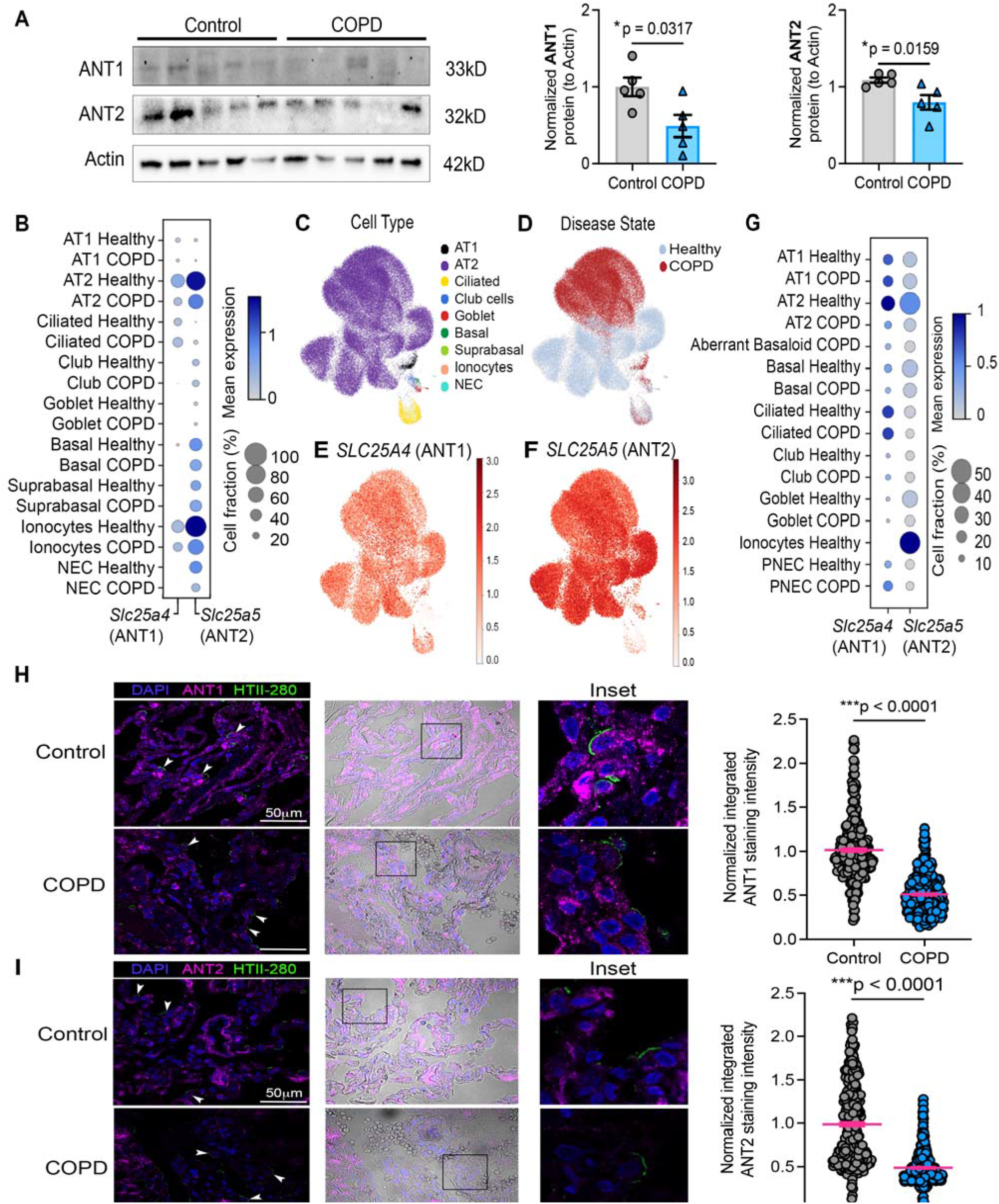
***SLC25A4* (ANT1) and *SLC25A5* (ANT2) are downregulated in COPD lung tissue and AT2 cells.** (A) Western blot analysis of ANT1 and ANT2 from whole human lung lysate from control and COPD patients, quantification on the right (n=5, each group). Data represent mean ± SEM. Statistics are calculated using the unpaired nonparametric Mann-Whitney test. (B) Single-cell RNA sequencing results obtained on isolated lung epithelial cells from normal control (n=4) and COPD (n=6) subjects (86). Mean expression and fraction of cells with gene expression of *SLC25A4* and *SLC25A5* for each cell cluster type are depicted. Circle size represents the percentage of cells expressing the gene, and color gradient represents the average expression in cells. (C) Uniform manifold approximation and projection (UMAP) visualization of annotated lung epithelial cell types. (D) UMAP visualization of disease state. (E) UMAP visualization of *SLC25A4* expression. (F) UMAP visualization of *SLC25A4* expression. (G) Single-cell RNA sequencing results obtained on normal control and COPD subjects (Sauler et al 2022). (H and I) Immunofluorescence of lung tissue from healthy control subjects and patients with COPD for ANT1 and ANT2, respectively. The white arrows indicate representative HTII-280+ AT2 cells (green). ANT (pink) staining intensity was quantified in AT2 cells across images (six to nine images per patient, n = 3 patients per group). Statistics are calculated using the unpaired nonparametric Mann-Whitney test. Scale bars, 50 μm (overview) and inset. Brightfield overlay images and scale bars are provided. *p < 0.05, **p < 0.01, ***p < 0.0001.

### Loss of ANT2 in lung epithelium and AT2 cells results in increased emphysema

AT2 cell function is important for normal alveolar homeostasis and the repair response to lung injury, such as with cigarette smoke (CS). To determine the role of ANT2 loss in lung injury and repair, we used the clinically relevant mouse CS model of emphysema/COPD. We generated a mouse with loss of Ant2 in all lung alveolar epithelial cells using the surfactant protein C (SPC) promoter (constitutively active, Sftpc-Cre^Blh^) (**Figure 2A**). We confirmed the loss of Ant2 expression in isolated AT2 cells at the gene and protein levels (**Figure S1A-C**). The constitutive Sftpc-Cre (Blh) line reflects Sftpc expression during lung development (43). We exposed 10- to 12-week-old mice to CS for 6 months. After completing 6 months of CS exposure, the Ant2-null^EPI^ mice develop exaggerated airspace enlargement (emphysema) (**Figure 2B-C**), as measured by alveolar cord length. Notably, under air condition, by 9 months of age, Ant2-null^EPI^ mice show a trend toward spontaneous emphysema with aging, similar to what is observed in humans with aging (44,45). The Ant2-null^EPI^ mice treated in air or smoke had a significant increase in bronchoalveolar lavage (BAL) macrophages compared to WT mice (**Figure 2D, S2A**). On pulmonary function testing, no difference in lung compliance and elastance was observed between WT and Ant2-null^EPI^ mice treated with CS after 6 months (**Figure S2B**) likely reflecting developmental compensation afforded by the Sftpc-Cre(Blh) allele’s broad embryonic recombination activity (43) allowing for the developing lung to compensate for Ant2 loss through lineage plasticity and developmental adaptation. These effects may be further obscured by the inherently stiffer lungs and attenuated remodeling response of the C57BL/6J strain (46,47).

**Figure 2:**
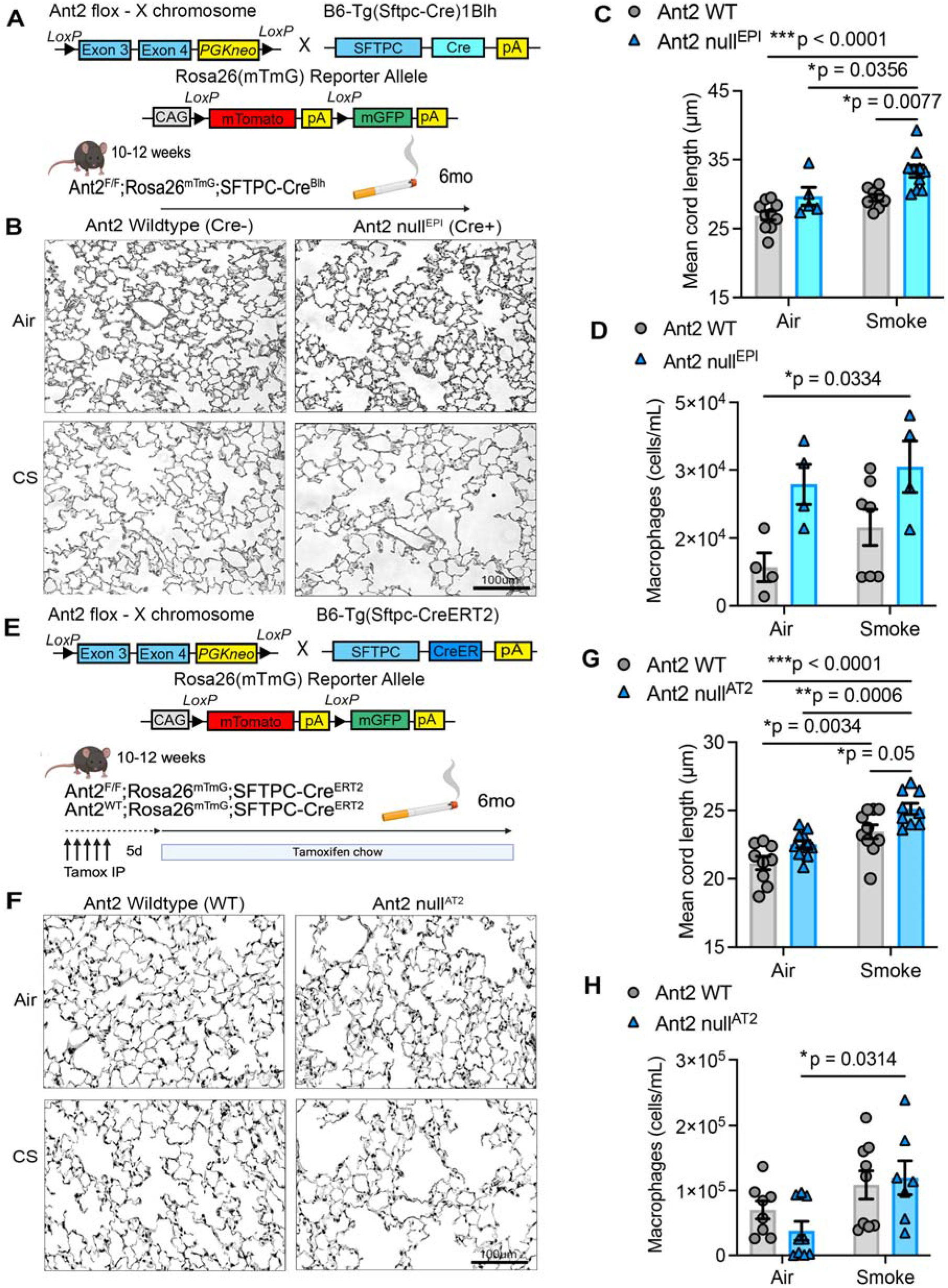
Loss of ANT2 in lung epithelial cells promotes a greater emphysema effect. (A) Schematic illustrating the cross-breeding of floxed *Ant2* mice with Sftpc-Cre mice to generate lung epithelial-specific Ant2 knockout mice. Outline of six-month smoking experiment timeline. (B) Representative black and white Gill’s stain images of paraffin-embedded lungs from wild-type (WT) or Ant2-null ^EPI^ mice treated with air or smoke (SM) for 6 months. (n = 5–10 mice per group). Scale bars, 100 μm. (C) Average alveolar chord length quantification from histological images. Each point represents a biologically independent mouse tissue sample, with average alveolar chord lengths measured from 5–10 mice per group, and 12–14 images per mouse. (D) Macrophage cell counts from BAL fluid. (E) Schematic illustrating the cross-breeding of floxed *Ant2* mice with Sftpc-Cre^ERT2^ mice to generate AT2 epithelial-specific Ant2 knockout mice. Outline of a two-month and a six-month smoking experiment timeline with mice given tamoxifen for 5 days prior to smoking. (F) Representative black and white Gill’s stain images of paraffin-embedded lungs from wild-type or Ant2-null ^AT2^ mice treated with air or smoke (SM) for 6 months. (n = 9-10 mice per group). Scale bars, 100 μm. (G) Average alveolar chord length quantification from histological images. Each point represents a biologically independent mouse tissue sample, with average alveolar chord lengths measured from 9-10 mice per group, and 12–14 images per mouse. (H) Macrophage cell counts from BAL fluid. Data are represented as mean ± SEM. *p < 0.05, **p < 0.01, ***p < 0.0001, with Statistics by two-way ANOVA with Tukey’s post hoc test. P values are noted.

To determine if loss of Ant2 specifically in AT2 cells in adult mice is sufficient to drive emphysema development, we generated an AT2 cell-specific knockout mouse using the Sftpc-Cre^ERT2^ mouse (**Figure 2E, S1D,E**) and exposed adult 10-12 weeks old mice to CS. Ant2-null^AT2^ mice developed enhanced emphysema after 6 months of CS exposure (**Figure 2F-G**). There as a significant increase in macrophages in the BAL in Ant2-null^AT2^ mice treated with CS compared to their air control group (**Figure 2H, S2C**), suggesting increased inflammatory response. Ant2-null^AT2^ mice treated with CS had reduced elastance and increased compliance on pulmonary function testing compared to the WT mice treated in air and CS, consistent with COPD patterns (**Figure S2D**). These results demonstrate that ANT2 expression in AT2 cells is essential for progenitor cell function and alveolar lung regeneration in response to cigarette smoke injury. Collectively, these findings establish that reduced ANT2 expression is associated with human COPD and that adult-specific loss of ANT2 in AT2 cells increases susceptibility to and progression of CS-induced emphysema. This suggests that loss of ANT2 is a mechanistic contributor to predisposing to poor alveolar repair and disease development.

### Loss of ANT2 alters mitochondrial morphology, oxygen use, and ATP production

To determine the impact of loss of ANT2 in mitochondrial structure and function, we first isolated AT2 cells from WT and Ant2-null^EPI^ mice and visualized the mitochondria using transmission electron microscopy (TEM) (**Figure 3A**). AT2 cells from Ant2-null^EPI^ mice exhibit an abnormal mitochondrial phenotype with a drastic absence of mitochondrial cristae and disrupted outer membranes (**Figure 3A**). There was no difference in the average mitochondrial area (**Figure S3A**). To analyze additional mitochondrial functional parameters, we generated ANT2 knockout (KO) cells using CRISPR-Cas9 in the Beas-2B human lung epithelial cell line. Loss of ANT2 protein was confirmed by Western blot analysis (**Figure S3B**). We determined mitochondrial membrane potential using tetramethyl rhodamine methyl ester (TMRM) and flow cytometry. Using FCCP as a positive control, we observed a significant decrease in membrane depolarization in ANT2 KO cells treated with CS extract (CSE) compared with control cells treated with CSE, control cells in normal media, and ANT2 KO cells in normal media. (**Figure 3B,C**). ANT2KO cells also showed a decrease in the NAD^+^/NADH ratio (**Figure 3D**) and total cellular ATP per cell (**Figure 3F**). To confirm metabolic dysfunction in the absence of ANT2, we performed the Seahorse MitoStress assay to assess cellular bioenergetics. ANT2 KO cells had significantly reduced basal oxygen consumption rate (OCR), proton leak, maximal and spare respiratory capacity compared to control cells (**Figure 3G**). We observed a significant decrease in ATP production rate from oxidative phosphorylation by Seahorse analysis (**Figure 3H**). We next confirmed the reduction in oxidative respiration by performing Seahorse analysis on murine AT2 cells isolated by FACS sorting for GFP+ cells from our WT and Ant2-null^AT2^ mice. In primary AT2 cells, loss of Ant2 resulted in a significant decrease in basal and ATP-linked respiration (**Figure 3I**). Consistent with the previously described role of ANTs in mitochondrial uncoupling, loss of Ant2 resulted in a reduction in proton leak which can contribute to excessive mitochondrial oxidative stress. These data demonstrate that loss of Ant2 results in abnormal mitochondrial function through structural changes and mitochondrial membrane depolarization, accompanied by decreased oxidative phosphorylation and ATP production. We next determined the transcriptional pathways perturbed by the loss of ANT2 in lung epithelial cells.

**Figure 3:**
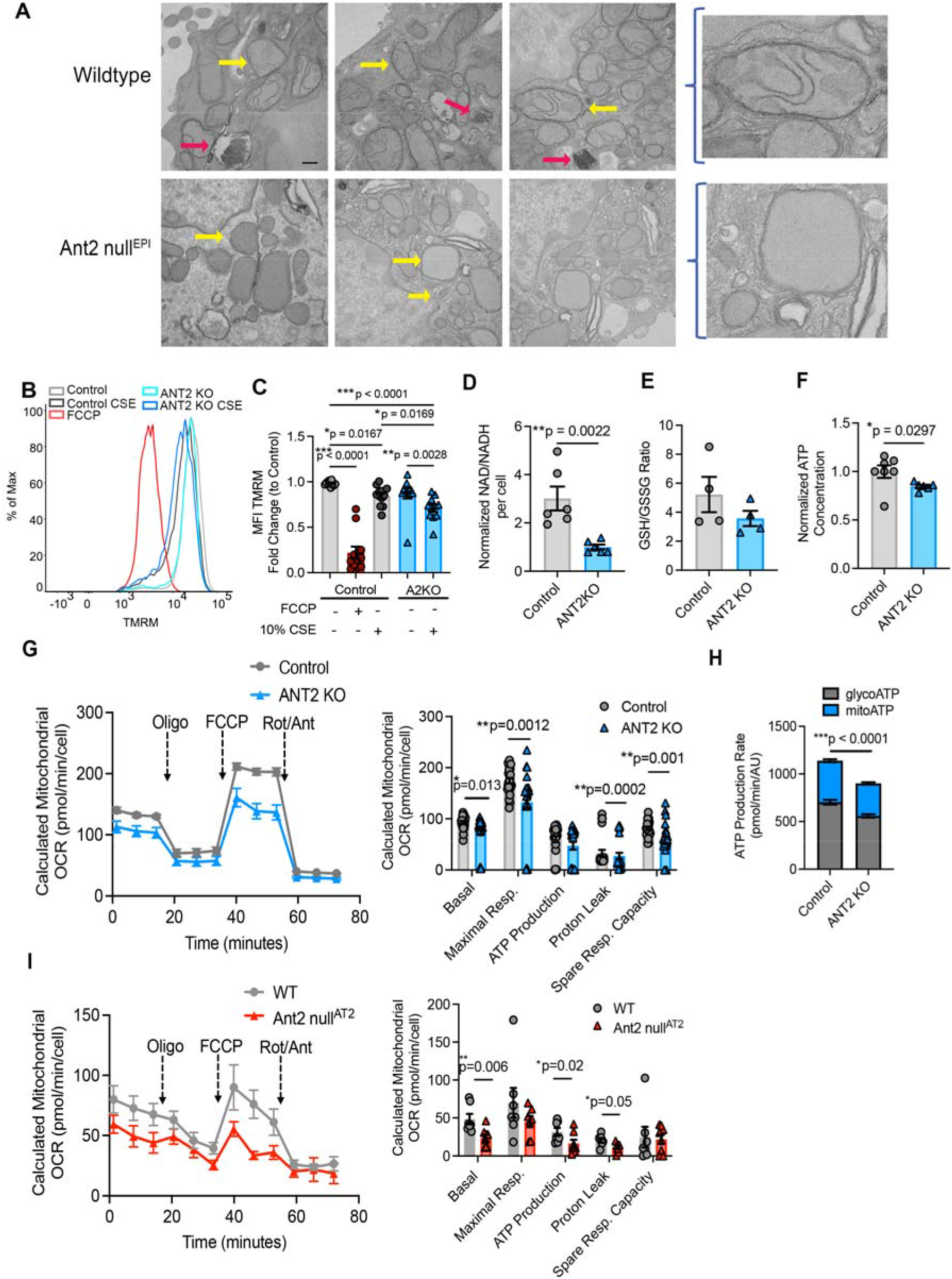
Loss of ANT2 alters mitochondrial function. (A) Transmission electron microscopy on isolated murine AT2 cells from wild-type or Ant2-null^EPI^ (n=2). Scale bar is 400 nm. Pink arrows indicate lamellar bodies found in AT2 cells and yellow arrows indicate mitochondria. (B) Histogram of flow cytometry of control and ANT2 KO cells treated with or without CSE for TMRM (ΔΨ m indicator) and analyzed in flow cytometry. (C) Bar graph shows the quantification of the mean TMRM fluorescence intensity across at least 3 experiments (quantified from 3 independent determinations). FCCP-treated groups served as a positive control. (D) Normalized NAD+:NADH ratios from control and ANT2 KO cell lysates (n=6 each group). (E) Normalized GSH/GSSG ratios from cell lysates (n=6 each group). (F) Normalized total cellular ATP from control and ANT2 KO cell lysates (n = 7 each group). (G) Seahorse Mito Stress Test was performed to measure oxygen consumption rate (OCR) on control and ANT2 KO cells (grey/blue; n = 24 per group) using Seahorse XF96. Cells were sequentially treated with DMEM, Oligo, FCCP, and rotenone/antimycin. Bar graphs quantify OCRs normalized to cell density, and Student’s t tests were performed to determine significance. P values are noted on the graphs. (H) ATP Rate Assay to measure total ATP and fractions of ATP produced from OXPHOS and glycolysis (blue/grey, respectively, n=46 per group). (I) Seahorse Mito Stress Test was performed to measure oxygen consumption rate (OCR) on isolated murine AT2 cells from WT and Ant2-null^AT2^ mice (grey/red; n = 8-9 per group) using Seahorse XF96. Cells were sequentially treated with DMEM, Oligo, FCCP, and rotenone/antimycin. Bar graphs quantify OCRs normalized to cell density, and Student’s t tests were performed to determine significance. P values are noted on the graphs. Data are represented as mean ± SEM. *p < 0.05, **p < 0.01, ***p < 0.0001, with Statistics by two-way ANOVA with Tukey’s post hoc test. P values are noted.

### ANT2 drives ROS production and oxidative stress

The mitochondria are important organelles for energy production, metabolic integration, and redox homeostasis. They are a key source of ROS (such as superoxide O_2_^•−^ and H_2_O_2_) that can act as signaling molecules and oxidize proteins, lipids, and DNA in the cell. We next asked whether loss of ANT2 alters oxidative stress in the lung epithelium. We stained mouse lung tissue from WT and Ant2-null^EPI^ mice for these oxidative modifications. Ant2-null mice had a significant increase in nitrotyrosine (NT; protein oxidation) and 4-hydroxynonenol (4-HNE, lipid peroxidation) at baseline in air-exposed animals compared to wild-type controls (**Figure 4A, B**). There were further increases in NT, 4-HNE, and 8-hydroxy-2-deoxyguanosine (8-OHDG, DNA damage) in Ant2-null mice after CS exposure for 6 months compared to CS-exposed WT mice (**Figure 4A-C**). The expression of antioxidants is an important cellular response to increased ROS levels that limits their negative effects (**Figure 4D**). We next evaluated bulk RNA sequencing data from AT2 cells isolated from air and CS-exposed Ant2-null and WT mice for superoxide dismutase (*Sod1*), *Sod2*, and catalase (*Cat*). There was a significant increase in both *Sod1* and *Sod2* in Ant2-null^EPI^ mice treated with CS, compared with WT (**Figure 4E**). We next confirmed that loss of ANT2 increases mitochondrial ROS, as assessed by MitoSox analysis, in both the presence and absence of CSE treatment, compared with control lung epithelial cells (**Figure 4F**). Finally, we found that ANT2KO cells exhibited exaggerated H_2_O_2_ production in the presence of CSE, compared with control cells, with no significant changes in peroxidase activity (**Figure 4G, H**). Increased H₂O₂ without changes in peroxidase activity indicates that oxidative stress primarily results from increased ROS production or weaker antioxidant defense, rather than decreased peroxidase function. We performed Gene Set Enrichment Analysis (GSEA) using the MitoCarta3.0 gene set, comprising over 1000 genes encoding the mitochondrial proteome (48), on bulk RNA sequencing data from AT2s isolated from CS-exposed Ant2-null and WT mice. Loss of Ant2 in the context of CS led to enrichment in detoxification and key metabolic pathways, including fatty acid oxidation (**Figure S3C**). With ANT2 loss, we observe increased oxidative stress, a compensatory rise in antioxidant enzymes, enhanced detoxification, ROS response, and fatty acid metabolism, along with elevated oxidative markers, indicating activation of an oxidative stress response linked to metabolic remodeling.

**Figure 4:**
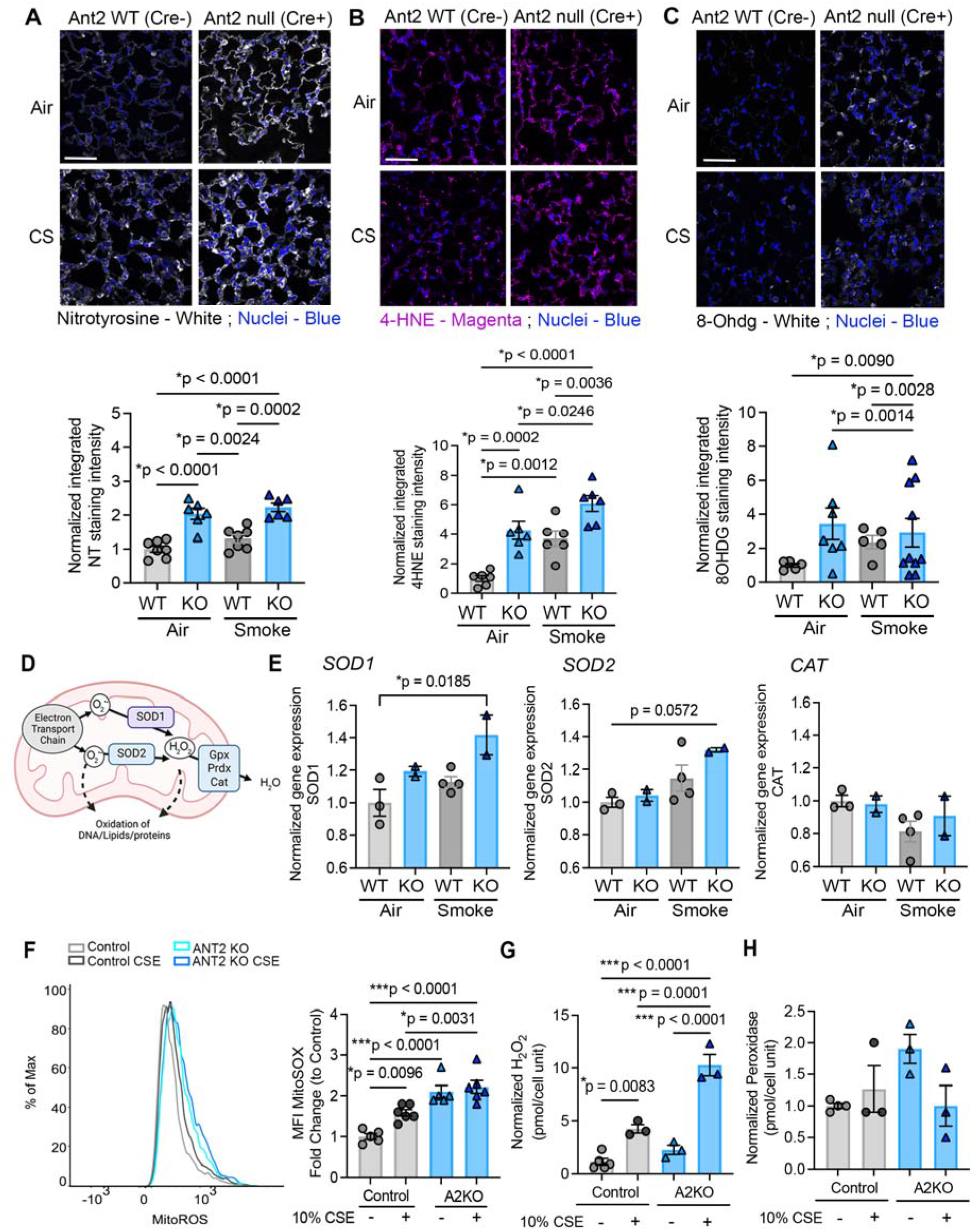
**Loss of ANT2 increases oxidative stress**. (A-C) Immunofluorescence of mouse lung tissue from wild-type and Ant2-null^EPI^ mice (seven to nine images per mouse, n = 3 mice per group). Nuclei are indicated in blue, nitrotyrosine in white, 4-HNE in magenta, and 8-OHDG in white. Quantification of the normalized fluorescence intensity of each respective oxidative stress marker. Scale bar is 100μm. (D) Schematic of antioxidant response to ROS generation in the mitochondria. (E) Normalized gene expression of Superoxide Dismutase 1 (*Sod1*), Superoxide Dismutase 2 (*Sod2*), and Catalase (*Cat*). (F) Histogram of flow cytometry of control and ANT2 KO cells treated with or without CSE for MitoSOX and analyzed in flow cytometry. The bar graph shows the mean MitoSOX fluorescence intensity across 3 experiments (n=5-6 per group). (G) H_2_O_2_ generation assessed in control and ANT2 KO cells with the Amplex Red probe/HRP read fluorescence with excitation at 530 nm and emission detection at 590 nm. (H) Peroxidase assay assessed in control and ANT2 KO cells, with fluorescence excitation at 530 nm and fluorescence emission at 590 nm. Statistics are calculated using the unpaired nonparametric Mann-Whitney test and two-way ANOVA with Tukey’s post hoc test when appropriate. Data are represented as mean ± SEM. *p < 0.05, **p < 0.01, ***p < 0.0001.

### Loss of ANT2 results in ferroptosis signatures and lipid peroxidation

Given that ANT2 loss impaired mitochondrial bioenergetics and increased oxidative stress in AT2 cells, we sought to define the downstream transcriptional consequences of ANT2 deficiency. We performed bulk RNA sequencing (bulk Seq) on Beas-2B cells with ANT2 knockout (KO) by CRISPR-Cas9, with or without CSE treatment. In ANT2 KO cells, these was enrichment in metabolic and oxidative stress, cellular stress adaptation pathways and cell fate pathways such as ferroptosis, compared to wild-type control cells (**Figure 5A**). We performed bulk RNA sequencing on isolated AT2 cells from WT and Ant2 null^EPI^ mice treated with CS for 6 months. There was significant enrichment in cellular stress, epithelial function, and metabolic signaling (**Figure S4A**). There was significant enrichment in ferroptosis in both ANT2KO cells treated with CSE and Ant2 null^EPI^ mice treated with CS compared to their WT counterparts, a distinct form of regulated cell death that is iron- and oxidative stress-dependent. GSEA analysis of the ferroptosis gene set demonstrates a normalized enrichment score (NES) of 1.45, p-value <0.0001, and FDR = 0.05 (**Figure 5B**). Additionally, in our bulk RNA-seq data from isolated murine AT2 cells from WT and Ant2 null^AT2^, some of the most upregulated genes included hemoglobin subunits (HBA/HBB) and antioxidant response genes (GPX3) (**Figure S4B**). The upregulation of hemoglobin subunits suggests altered iron handling or redox balance, further linking the phenotype to ferroptosis biology. Examining the differentially expressed genes within the ferroptosis gene set from Control CSE and ANT2 KO CSE cells, a decline is observed in genes involved in the glutathione/antioxidant pathway of ferroptosis, alongside an increase in genes related to the iron and lipid pathways of ferroptosis (**Figure 5C**). This transcriptional profile aligns with disrupted redox balance and the activation of compensatory mechanisms in iron management and membrane lipid remodeling, characteristics associated with susceptibility to ferroptosis. Ferroptosis has been implicated in COPD pathogenesis, with most studies focused on macrophages and lung epithelial cells broadly (33,34). However, the relationship between ferroptosis and mitochondrial function in AT2 cells remains largely unexplored, with prior work primarily conducted in other lung injury models (49–52). We next sought to directly examine how mitochondrial dysfunction induced by loss of ANT2 contributes to ferroptotic cell death in AT2 cells in COPD. Analyzing scRNA sequencing data from healthy and COPD subjects, UMAP from ferroptosis-related genes showed partial separation of COPD and healthy lung epithelial cells, indicating disease-related changes in ferroptosis transcription in lung epithelial cells (**Figure 5D**). Gene expression in COPD AT2 cells shows suppressed iron and antioxidant defenses (*HMOX1, SLC7A11, GPX4*), lipid remodeling enzymes (*LPCAT3, ACSL4*), and increased ferritinophagy (*NCOA4*). (**Figure S4C**). This indicates exhausted antioxidant response and selective iron release. We measured several key ferroptosis-related proteins and found distinct changes in ANT2KO samples treated with CSE compared to controls. There was a combined pattern of decreased GPX4, increased NCOA4, HMOX1, and FTH *in vitro* (**Figure 5E**), all suggesting activation of an iron-dependent lipid peroxidation process typical of ferroptosis. However, in the lung lysate from wild-type and ANT2-null^AT2^ mice treated in air or CS for 6 months, the Ant2-null^AT2^ mice had increased Ncoa4 and Hmox1 protein but an increase in Slc7a11 and Gpx4, indicating iron stress with a mounting compensatory antioxidant response to the stress (**Figure S4E**). To further assess ferroptosis, flow cytometry was conducted to quantify lipid peroxidation in both control and ANT2 knockout cells, with and without CSE stimulation. RSL3 served as a positive control for ferroptosis induction, demonstrating approximately a 3.5-fold increase in lipid peroxidation (**Figure 5F**). Treatment with a ferroptosis inhibitor, such as liproxstatin, significantly reduced lipid peroxidation. Importantly, ANT2 KO cells showed a marked increase in baseline lipid peroxidation (about 2.5 times higher) even without CSE stimulation which was significantly reduced by liproxstatin. However, when assessing cell membrane integrity, only the ANT2KO cells treated with CSE exhibited an increase in dead cells with damaged membranes. (**Figure 5G**). Additionally, the total 4-HNE from lung lysate increased in the Ant2-null^AT2^ mice treated in air or CS for 6 months compared to their WT counterparts (**Figure S4D**). These data support that loss of ANT2 promotes susceptibility to ferroptosis-related oxidative lipid damage and cell death. However, ANT2KO treated with CSE showed increased lipid peroxidation that was not significantly prevented by ferroptosis inhibitors (**Figure 5G**), implying that the stimulus triggers cell death via ferroptosis-independent or mixed pathways. Ferroptosis has several hallmarks including oxidative stress, iron availability, and lipid peroxidation. We next determined how loss of ANT2 impacts these requirements for ferroptosis.

**Figure 5:**
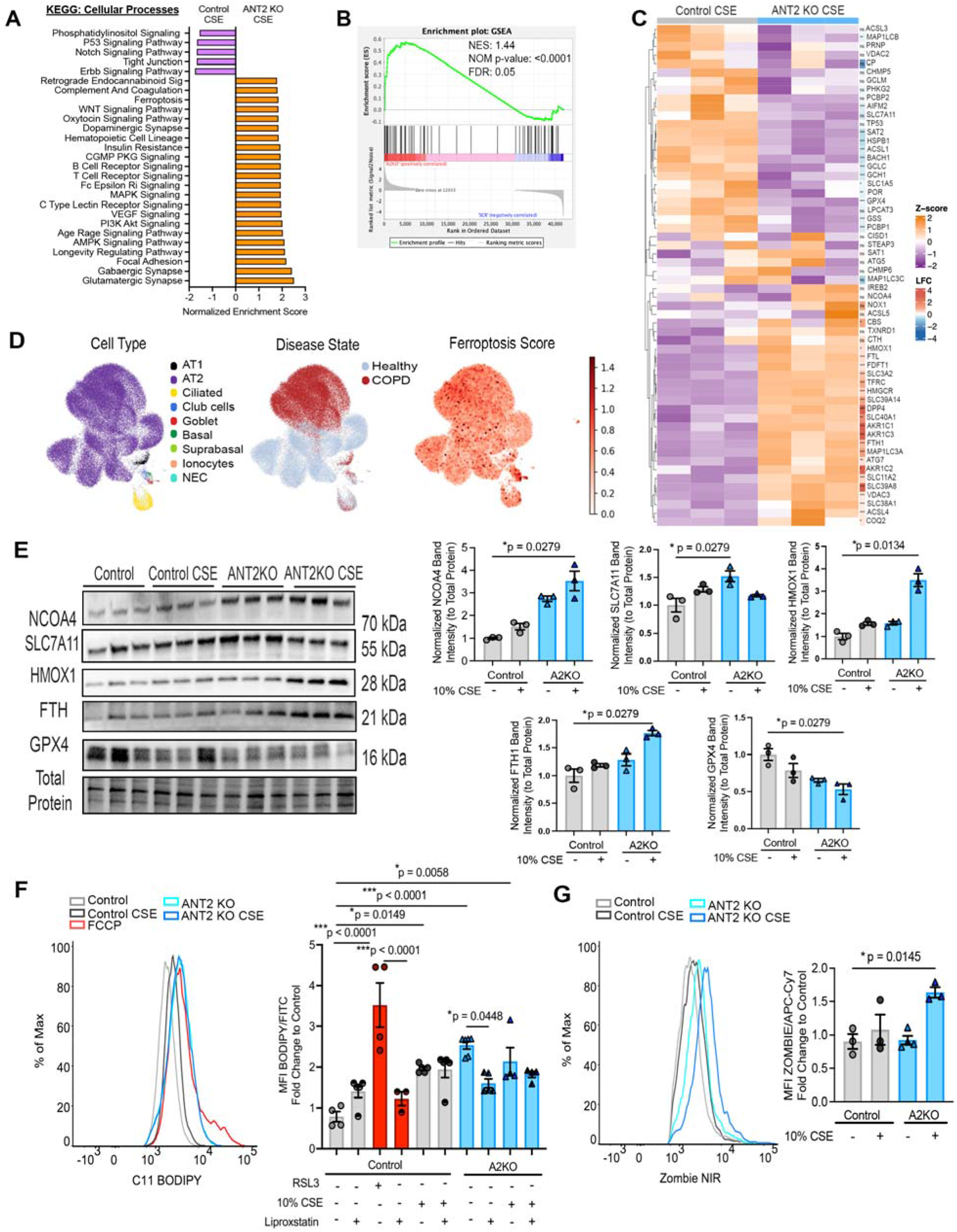
**Loss of ANT2 show enrichment of ferroptosis pathways**. (A) Gene set enrichment analysis (GSEA) pathway analysis matched to Kyoto Encyclopedia of Genes and Genomes enrichment category for Cellular Processes was performed on significantly differentially expressed gene lists (DEGs) generated comparing CRISPR-Cas9 WT and ANT2 KO Beas2b cells treated with 10% CSE (n =3 per treatment group). Significantly enriched pathways are indicated by (p-value < 0.05 and FDR < 0.05). (B) Enrichment plot for Ferroptosis pathway (WP_Ferroptosis; GSEA Msigdb) with Normalized enrichment scores (NES), p-value, and FDR values. (C) DEGs for the Ferroptosis Pathway; Z-score, log2(Fold Change), and Condition are indicated in the legend. (D) UMAPs showing ferroptosis score across cell types from isolated lung epithelial cells from normal control (n=4) and COPD (n=6) subjects. (E) Western blot for ferroptosis genes performed on cell lysates from CRISPR-Cas9 WT and ANT2 KO Beas2b cells either treated with culture media or 10% CSE. Respective quantifications (F) Histogram of flow cytometry of control and ANT2 KO Beas2b cells treated with or without CSE for C11-BODIPY for lipid peroxidation and analyzed in flow cytometry. The bar graph shows the mean C11-BODIPY fluorescence intensity across 3 independent experiments (n=3-6 per group). RSL3, a ferroptosis inducer, served as a positive control. Liproxstatin-1 is a ferroptosis inhibitor. (G) Histogram of flow cytometry of control and ANT2 KO Beas2b cells treated with or without CSE for ZOMBIE NIR for viability and analyzed in flow cytometry. The bar graph shows the mean ZOMBIE NIR fluorescence intensity across 3 independent experiments (n=3 per group). (G) Statistics are calculated using the unpaired nonparametric Mann-Whitney test and two-way ANOVA with Tukey’s post hoc test when appropriate. Data are represented as mean ± SEM. *p < 0.05, **p < 0.01, ***p < 0.0001.

### Loss of ANT2 promotes iron accumulation through Mitoferrin-1

Previous studies show iron accumulation in COPD lungs, indicating disrupted iron balance in disease. Perls’ staining found minimal iron in normal lungs, but increased iron in COPD lungs (**Figure S5A**). Excess free iron is an important catalyst for ferroptosis, as it drives Fenton reactions (the conversion of hydrogen peroxide) to produce highly reactive intracellular ROS that alter lipid membranes. To elucidate the mechanistic basis of iron homeostasis in ANT2-driven ferroptosis, we next measured intracellular iron levels in ANT2KO cells. We observe increased cellular and mitochondrial iron in the ANT2 KO cells at baseline compared to the WT cells (**Figure 6A,B**). CSE caused further increases specifically in mitochondrial iron in ANT2 KO cells treated with CSE compared to WT cells (**Figure 6B**). We confirmed the increase in iron with loss of Ant2 in mice, demonstrating an increase in total cellular iron in lung tissue from Ant2-null^AT2^ mice treated with air or CS for 6 months compared to WT mice (**Figure 6C**). Additionally, we confirmed increased heme in Ant2-null^AT2^ mice, as total heme was significantly higher in Ant2-null^AT2^ mice treated with air or CS for 6 months compared to WT mice (**Figure 6D**). To elucidate the dynamics of iron transport and distribution, we analyzed gene expression related to iron import, export, and utilization by comparing ANT2 and wild-type (WT) specimens. Our findings revealed upregulation of iron import genes such as *SLC25A37* (Mitoferrin-1, MFRN-1), Transferrin Receptor (*TFRC*), and *SLC39A14* (ZIP14), alongside downregulation of *SLC25A28* (Mitoferrin-2, MFRN-2) (**Figure 6E**). No significant differences were observed in the expression levels of *SLC11A2* (DMT1) and the iron exporter *SLC40A1* (Ferroportin, FPN). Notably, particular interest was directed toward MFRN-1, which is predominantly expressed in erythroid lineages and functions as a regulator that determines mitochondrial iron utilization to support cellular growth and metabolism, rather than merely governing the quantity of iron entering the cell. We detected increased MFRN1 in AT2 cells from Ant2-null^AT2^ mice compared to WT mice by IF staining and western blot analysis of lung tissue (**Figure 6G, S5E**). When MFRN1 is knocked down in ANT2KO cells, an increase in TFRC and ferroportin levels was observed, while MFRN-2 levels remain unchanged relative to control cells (**Figure 6F**). ANT2KO cells had reduced cellular proliferation compared to WT cells, with further reduction in proliferation with knockdown of MFRN1 (**Figure 6H**); however, ANT2KO cells at baseline exhibit increased heme content but no decrease in heme or cellular and mitochondrial iron with knockdown of MFRN1 (**Figure 6I, S5B-D**). Elevated TFRC expression may signify activation of iron-responsive signaling pathways, whereas increased ferroportin could represent a compensatory mechanism to limit excess cytosolic iron or heme-associated stress. To assess the potential of iron chelation to modulate lipid peroxidation in the context of loss of ANT2, we performed flow cytometry experiments in control and ANT2KO cells treated with or without CSE. ANT2KO at baseline showed elevated lipid peroxidation, which was abrogated by Deferoxamine (DFO), a known iron chelator (**Figure 6J**). In this case, ANT2KO treated with CSE showed reduced lipid peroxidation when co-treated with an iron chelator, compared with Liproxstatin-1 treatment (**Figure 5I**). Collectively, these key findings demonstrate that Mitoferrin-1 is essential for mitochondrial iron uptake and ferroptosis susceptibility specifically in the context of ANT2 deficiency, driven by labile iron accumulation, revealing a previously unrecognized link between mitochondrial metabolic state, ANT2 and iron-dependent cell death.

**Figure 6:**
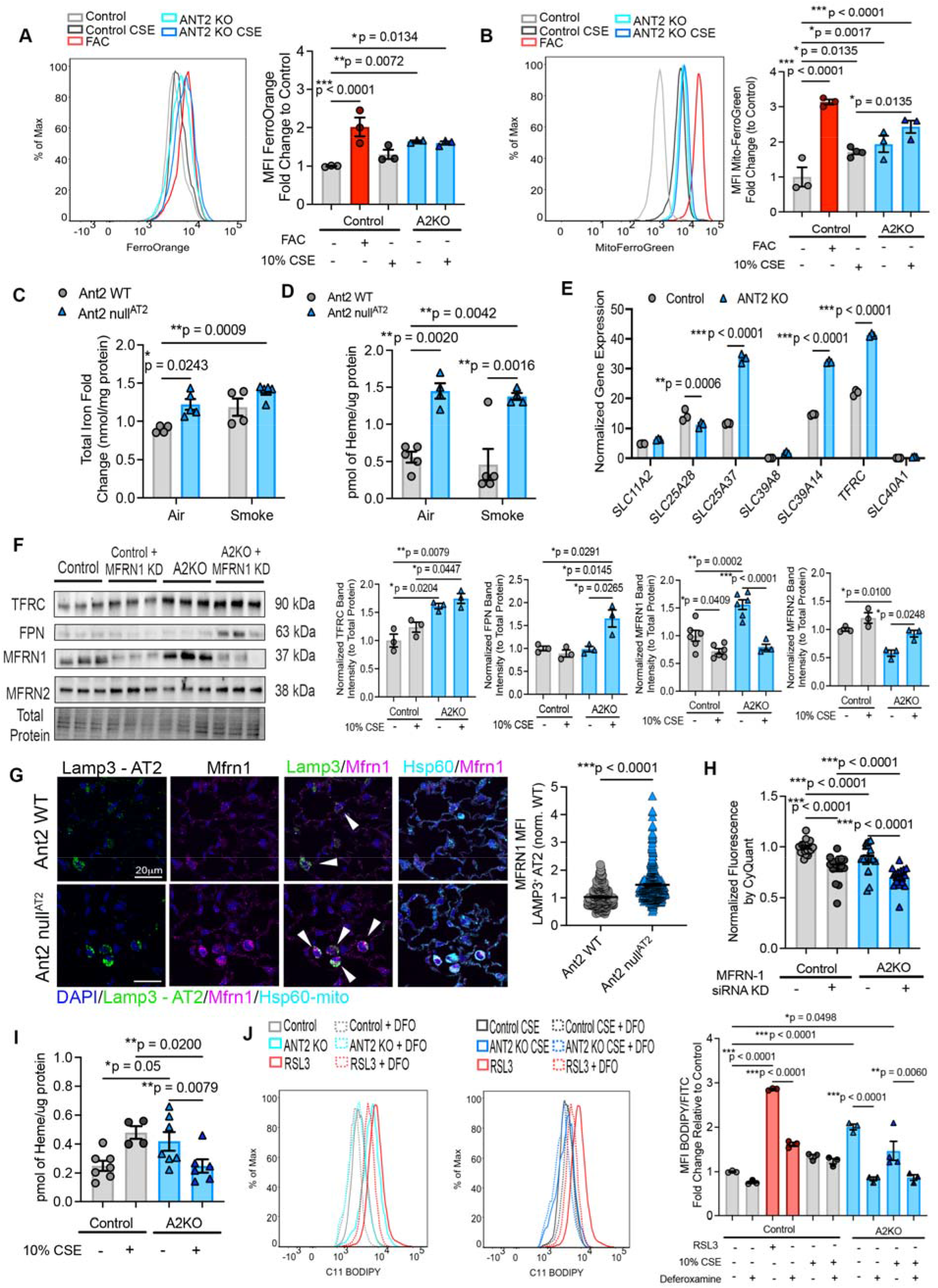
Loss of ANT2 alters iron homeostasis, including through Mitoferrin-1 (Mfrn1). (A) Histogram of flow cytometry of control and ANT2 KO cells treated with or without CSE for FerroOrange (intracellular iron) and (B) MitoFerroGreen (mitochondrial iron) analyzed in flow cytometry. The bar graph shows the mean probe fluorescence intensity across at least 3 experiments (derived from 3 independent determinations). Ferric Ammonium Citrate (FAC)-treated groups served as a positive control. (C) Iron assay quantification of WT and Ant2-null ^AT2^ mice treated with or without cigarette smoke for 6 months detecting total iron. (D) Heme assay quantification in WT and Ant2-null ^AT2^ mice treated with or without cigarette smoke for 6 months. (E) Normalized Gene expression of iron import and export genes. *SLC11A2* (DMT1), *SLC25A28* (Mitoferrin2), *SLC25A37* (Mitoferin1), *SLC39A8* (ZIP8), *SLC39A14* (ZIP 14), *TFRC*, *SLC40A1* (Ferroportin). Statistics are calculated using the unpaired nonparametric Mann-Whitney test. (F) Western blot for iron import and export genes performed on cell lysates from CRISPR-Cas9 control ad ANT2 KO Beas2b cells either treated with culture media or 10% CSE. Respective quantifications. (G) Immunofluorescence of mouse lung tissue from WT and Ant2 null^AT2^ mice for Mitoferrin-1 (magenta), LAMP3 (green) for AT2 cells, and HSP60 (cyan) for mitochondria. The white arrows indicate representative LAMP3+ AT2 cells (green). Mitoferrin-1 staining intensity was quantified in AT2 cells across images (6-9 images per mouse, n = 3 mice per group). Scale bar is 20μm. (H) Cellular proliferation was assessed using the CyQUANT Cell Proliferation Assay. Fluorescence intensity, which is proportional to cellular nucleic acid content, from CRISPR-Cas9 control and ANT2 KO Beas2b cells either treated with or without siRNA knockdown of Mitoferrin-1. (I) Heme content was determined by oxalic acid-induced conversion of heme to fluorescent protoporphyrin IX (PPIX) for CRISPR-Cas9 control and ANT2 KO Beas2b cells treated with culture media or 10% CSE for 24 hrs. Background-corrected fluorescence (heated minus unheated samples) was normalized to total protein concentration. (J) Histogram of flow cytometry of control and ANT2 KO Beas2b cells treated with or without CSE for C11-BODIPY for lipid peroxidation and analyzed by flow cytometry. The bar graph shows the mean C11-BODIPY fluorescence intensity across 3 independent experiments (n=3-6 per group). RSL3, a ferroptosis inducer, served as a positive control. Deferoxamine (DFO) is an iron chelator. Statistics are calculated using the unpaired nonparametric Mann-Whitney test and two-way ANOVA with Tukey’s post hoc test when appropriate. Data are represented as mean ± SEM. *p < 0.05, **p < 0.01, ***p < 0.0001.

### ANT2 regulates AT2 cell regenerative capacity and prevents emphysema development

Given that ferroptotic death of AT2 cells would likely deplete the alveolar progenitor pool, we next examined whether ANT2 loss independently impairs epithelial repair capacity. We generated 3D alveolar organoids using primary mouse AT2 cells from wild-type and ANT2-null mouse lungs. AT2 cells with loss of Ant2 have significantly lower organoid colony-forming efficiency and size (**Figure 7A**). Given the role of AT2 in human COPD and evidence of functional impact from loss-of-function studies, we determined whether therapeutic reconstitution of AT2 expression could restore AT2 cell plasticity and prevent the development of emphysema. We quantified the number of GFP+SPC+ AT2 cells from WT and Ant2 null^AT2^ mice and found a significant reduction in the number of AT2 cells in the Ant2 null^AT2^ mice (**Figure 7B**). We generated a new transgenic mouse model allowing for tissue-specific and temporal control of ANT2 expression using a Cre-recombinase and tetracycline-inducible system (**Figure 7C, S6A**). We crossed this mouse with the Sftpc-Cre^ERT2^ mouse to enable AT2-cell-specific Ant2 expression. In isolated AT2 cells, there was a 2-fold increase in *Slc25a5* (Ant2) gene expression with no change in *Slc25a4* (Ant1) (**Figure 7E, S6B-C**). In the 3D alveolar organoid assay, overexpression of ANT2 restores organoid growth and increases colony-forming efficiency, and increases the number of AT2 cells in the Ant2 Oexp^AT2^ mice (**Figure 7D, S6D**). Overexpression of ANT2 resulted in decreased 4-HNE staining following 6 months of CS exposure, indicating reduced lipid peroxidation and suggesting attenuated ferroptosis (**Figure 7G**). Overexpression of ANT2 resulted in decreased iron and heme content at baseline (**Figure S6E-F**), together indicating reduced oxidative stress and iron overloading. We next crossed Ant2-null mice with the Ant2-Overexpressing transgenic mouse (**Figure 7J**) to test whether reconstitution of Ant2 would restore AT2 cell progenitor capacity. Similarly, reconstitution of Ant2 in AT2 cells restored organoid growth and size (**Figure 7K**) compared to AT2 cells from Ant2 KO mice. Analysis of precision-cut lung slices (PCLS) from mice with AT2-cell-specific overexpression of ANT2 shows increased tissue metabolism by WST-8 assay and higher total ATP compared to PCLS from WT mouse lungs (**Figure 7H, I**). To test if increased Ant2 expression in AT2 cells alters the development of emphysema, we exposed mice with AT2-cell-specific Ant2 overexpression (Ant2^OExp_AT2^) or WT Ant2 to CS for 6 months. The Ant2^OExp_AT2^ mice were protected from emphysema development compared to WT mice exposed to CS (**Figure 7F**). Together, these data demonstrate a previously unknown role for ANT2 in regulating AT2 stem cell function, in susceptibility to iron-import-mediated ferroptosis via mitoferrin-1, and in its key role as a driver of lung repair and emphysema development.

**Figure 7:**
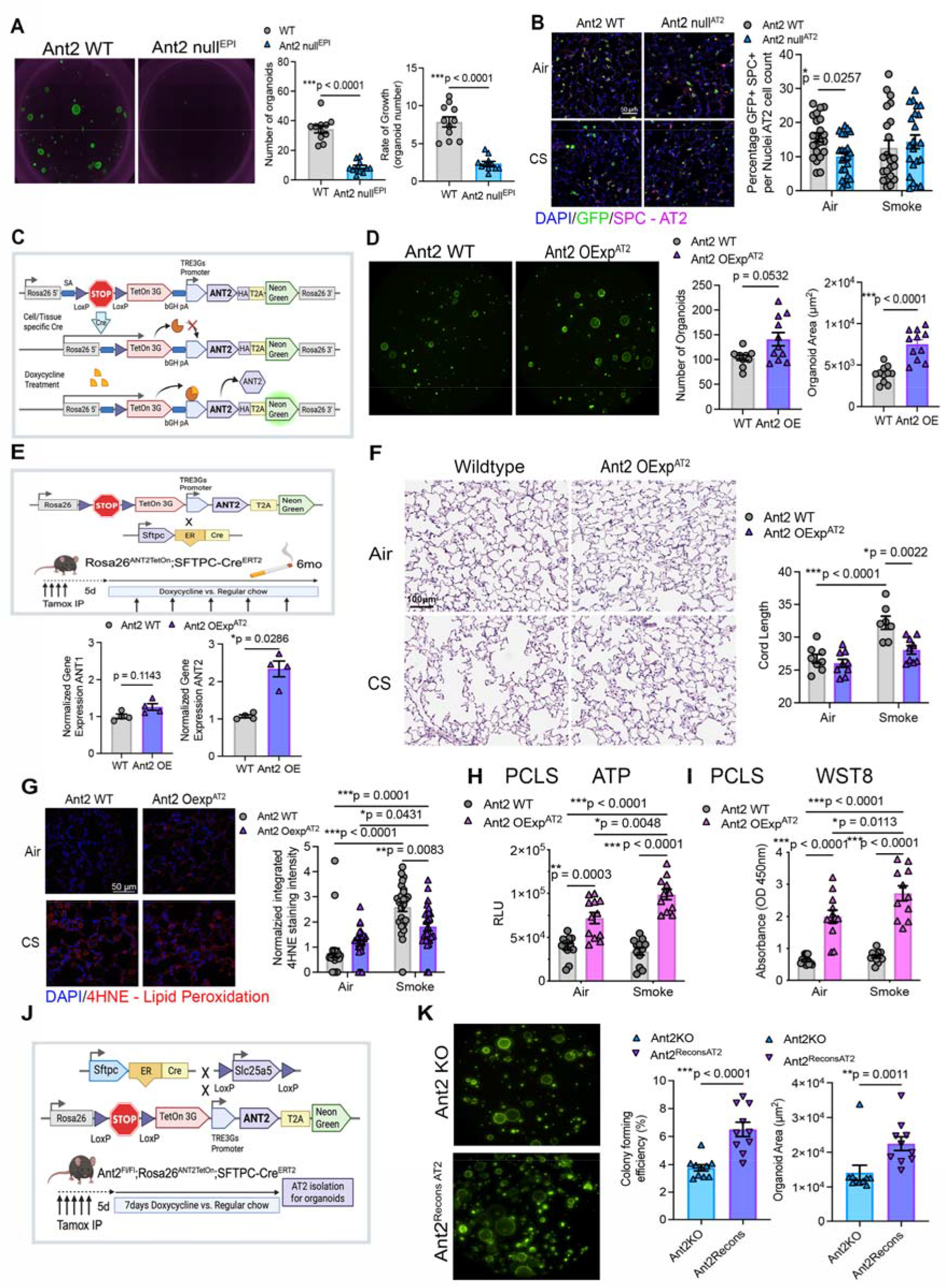
Overexpression or Reconstitution of ANT2 recovers AT2 progenitor cell viability and proliferative capacity. (A) AT2 cell-derived 3D alveolar organoids from wild-type or Ant2-null^EPI^ mice. Respective quantification of the number of organoids on day 14 and the rate of growth. Cell isolation from n=3 mice per group. Data is representative of 3 independent experiments. (B) Immunofluorescence of wild-type or Ant2-null ^AT2^ mice lungs treated with air or smoke (SM) for 6 months. (n = 3 mice per group). DAPI indicated in blue, GFP indicated in green, and SPC indicated in red. Respective quantification of GFP+SPC+ cells across the entire lung. Scale bar is 50μm. (C) Schematic of Ant2^OExp_AT2^ mice with tissue-specific and temporal control of ANT2 expression using a Cre-recombinase and tetracycline-inducible system with treatment of doxycycline. See the methods for additional details. (D) 3D alveolar organoids from wild-type or Ant2^OExp_AT2^ mice. Quantification of the number of organoids and average organoid area. (E) Schematic of the cross of transgenic mouse lines to generate AT2 cell-specific overexpression of ANT2 (Ant2^OExp_AT2^) with an outline of the six-month smoking experiment timeline. (F) Average alveolar chord length quantification from histological images of WT and Ant2^OExp_AT2^ mice treated with air and cigarette smoke for 6 months. Each point represents a biologically independent mouse tissue sample, with average alveolar chord lengths measured from 7–10 mice per group, and 12–14 images per mouse. Scale bar is 100μm. (G) Immunofluorescence of wild-type or Ant2^OExp_AT2^ mice lungs treated with air or smoke (SM) for 6 months. (n = 3 mice per group). DAPI is indicated in blue, and 4HNE is indicated in red. Quantification of the normalized fluorescence intensity of 4HNE oxidative stress marker. (H-I) PCLS from wild-type or Ant2^OExp_AT2^ mice lungs treated with or without 10% CSE for 24 hours followed by (H) ATP assay or (I) WST8 assay. n=11-12 PCLS per treatment. Data is representative of two independent experiments. (J) Schematic of Ant2^Recon_AT2^ mice for AT2 isolation and organoid analysis. (K) 3D alveolar organoids from Ant2 null^AT2^ or Ant2^Recon_AT2^ mice. Quantification of the number of organoids and average organoid area. All data are represented as mean ± SEM. Statistics are calculated using the unpaired nonparametric Mann-Whitney test and two-way ANOVA with Tukey’s post hoc test when appropriate. *p < 0.05, **p < 0.01, ***p < 0.0001.

## Discussion

The ability of the lung epithelium to repair after injury is an essential and metabolically demanding process for lung homeostasis. COPD is characterized by progressive epithelial dysfunction, abnormal AT2 cell behavior, and loss of reparative capacity, which all contribute to lung destruction. Although mitochondrial dysfunction has been implicated in impaired lung repair (5,53), the key mitochondrial factors governing this process remain unclear. Here, we identify a novel role for ANT2 as an important regulator of AT2 progenitor cell function, ferroptosis, and alveolar repair. Analysis of human COPD lung tissue reveals that ANT2 expression is reduced at all disease stages, suggesting that ANT2 deficiency is sustained across both early and late stages of disease development and offers an opportunity for therapeutic intervention early in pathogenesis before substantial lung destruction.

We delineated the function of ANT2 in AT2 progenitor cells and alveolar repair using human COPD tissue and cigarette smoke (CS) exposure models. Our findings demonstrate that ANT2 plays a pivotal role in maintaining metabolic equilibrium, regulating iron homeostasis, and modulating AT2 cell fate and regenerative capacity. Single-cell RNA sequencing confirmed ANT2 enrichment in AT2 cells, which have high proliferative capacity. Loss of ANT2 in a murine COPD model exacerbated emphysema with loss of AT2 cells, increased bronchoalveolar lavage macrophage counts, and worsened pulmonary function. Research on mitochondrial gene modulation in COPD remains limited. Existing mouse studies have demonstrated that deficiency in mitophagy-related *Pink1* can protect against CS-induced emphysema (54) and *Fxn*/*Sco2* protecting against mucociliary impairment in the airway (35). Deletion of mitochondrial complex I component *Ndufs4* during lung development was shown to activate AT2 cell stress responses which drive abnormal differentiation and derail alveolar structure during lung development (55). Our study is the first to demonstrate that ANT2 deficiency in AT2 cells impairs their progenitor stem cell function and contributes to abnormal alveolar repair and emphysema progression in the adult lung.

Four ANT isoforms exist with both ANT1 and ANT2 being expressed in human lung tissue and influence lung epithelial cell function (39). Whether a single isoform predominantly governs AT2 stem cell fate has not been established. Our prior studies showed that global ANT1 deletion protects against CS-induced emphysema by limiting monocyte-derived macrophage recruitment and alveolar destruction (56), and that ANT2 provides greater protection than ANT1 against CS injury in ciliated airway cells (39). The present study now establishes a specific and detrimental role for ANT2 deficiency in AT2 cell progenitor function and COPD progression.

Mechanistically, ANT2 loss in AT2 cells reduces ATP production and oxygen consumption, elevates ROS, and causes mitochondrial iron overload via mitoferrin-1, collectively promoting ferroptosis. ANT2 thus functions as both a sensor and amplifier of oxidative stress and iron imbalance upstream of ferroptosis-mediated cell death. Cells with loss of ANT2 had evidence of compensatory antioxidant responses including upregulation of *SOD1*, *SOD2*, and glutathione-related *SLC7A11*. However, these protective responses are insufficient to counterbalance the ROS and excessive iron-driven state. This underscores the vulnerability of ANT2-deficient cells to oxidative stimuli and the detrimental effects on progenitor cell function. Features of ferroptosis have been observed in AT2 cells in acute lung injury (ALI) models (49,51,52,57) and disrupted iron homeostasis is associated with inflammation, oxidative stress, and COPD-like pathology including airspace enlargement and airway thickening (33,58). We found that ANT2-deficient cells, as observed in the lungs of COPD patients, have increased susceptibility to ferroptosis which could be reverse by iron chelation and ANT2 expression. This suggests that ANT2 and mitochondrial regulation of ferroptosis may provide an early intervention point and therapeutic target to improve AT2 cell survival.

ANT2 deficiency also alters mitochondrial iron transport through increased Mitoferrin-1 (MFRN1/SLC25A37) expression while decreasing Mitoferrin-2 (MFRN2/SLC25A28). These mitochondrial iron importers are essential for heme and Fe-S cluster synthesis and are reciprocally regulated. IRP1/2 stabilizes MFRN1 mRNA under low-iron conditions while promoting MFRN2 degradation via FBXL5 (59). MFRN1 is predominant in erythroid cells with high iron demands; MFRN2 is the primary isoform in non-erythroid cells, where its silencing triggers compensatory MFRN1 upregulation to sustain heme synthesis (59). MFRN1 half-life is prolonged during proliferation or stress, unlike MFRN2 (60,61). In our data, increased MFRN1 expression appears to be a compensatory shift necessary for survival, given that knockdown of MFRN1 in ANT2-null cells decreases cellular proliferation. This compensatory response in MFRN1 promotes a state of excessive iron that establishes ferroptosis sensitivity. This gives more insight into the relationship between ANTs and mitoferrins. ANTs have been previously implicated in heme biosynthesis (62,63) and possibly form part of the mitochondrial heme metabolism complex (64), but mostly in yeast or erythroid cell line models. Consistent with a role in heme regulation, ANT2 loss is associated with elevated heme levels and increased HO-1 expression, the enzyme responsible for heme degradation and iron release, which may further contribute to cellular iron overload. The precise relationship between MFRN isoform dysregulation, iron overload, and ferroptosis in AT2 cells warrants further investigation.

These findings have therapeutic implications. Iron chelation reduces emphysema severity in preclinical models (35), and COPD patients frequently exhibit lung iron overload alongside systemic iron deficiency linked to exacerbations (65). The therapeutic use of iron chelators is in part hindered by systemic side-effects and impairment in other essential pathways. In our ANT2 KO model, iron chelation with deferoxamine improved organoid growth, supporting that iron regulation may be a viable therapeutic strategy. Importantly, beyond iron chelation, ANT2 overexpression diminishes emphysema development and restores AT2 progenitor capacity in mice, establishing ANT2 as a promising therapeutic target. Preclinical ANT2 research has largely focused on cancer immunotherapy (66,67), metabolic disease (68), and wound healing (69). Our work extends the relevance of ANT2 to lung disease. In addition to its canonical role as an ADP/ATP translocase, ANT2 participates in the mitochondrial permeability transition pore complex (70), functions as a mitochondrial uncoupler, and serves as an RNA translocon (71), highlighting the breadth of pathways that may be disrupted by ANT2 deficiency in COPD.

Several limitations and open questions remain. This study focuses on ANT2 in AT2 cells. The contributions of ANT1 and other isoforms in AT2 cell biology are important subjects for future work. Our prior studies show that global ANT1 deficiency is protective against macrophage-driven emphysema (56) and associated with airway remodeling in aging lungs (72). Additional AT2 cell functions are not examined here such as differentiation to AT1 cells, lipid regulation, and surfactant production, which are also important in lung homeostasis. Other progenitor cells such as basal cells reside in the airway epithelium, providing another important cell population to study the role of ANT2. AT2 cell heterogeneity is another key consideration as distinct subclusters may exhibit differential ANT2 expression depending on disease state (73). Spatiotemporal multi-omics could provide informative granular data on ANTs and AT2 diversity during repair. Finally, most human AT2 cell samples derive from severe or end-stage disease. iPSC-derived AT2 models would enable earlier-stage mechanistic studies and are an important future direction.

In summary, ANT2 loss in AT2 cells promotes ferroptosis, impairs lung progenitor self-renewal, and drives emphysema progression in COPD. Loss of ANT2 leaves AT2 cells with insufficient energy for regeneration and results in increased oxidative stress, iron overload, and heightened susceptibility to cell death by ferroptosis. Collectively, this reduces the functional AT2 progenitor pool and repair capacity of the lung. While we have demonstrated the importance of ANT2 in COPD, this work has broad important implication in other lung disease including aging, pulmonary fibrosis and acute lung injury. Therapeutic restoration of ANT2 represents a promising strategy. Broader approaches to AT2-targeted lung repair, including AAV or lipid nanoparticle-mediated gene therapy and mitochondrial transfer via stem cells or engineered nanoparticles (74–77), may complement ANT2-directed interventions. We propose that optimizing AT2 cell metabolic state through ANT2 or other mitochondrial targets at early disease stages would positively impact lung repair and prevent emphysema progression. These studies are the first to define how ANT2-mediated mitochondrial dysfunction drives ferroptosis in COPD, with broader applicability to lung aging and other lung diseases. Targeting ANT2 to limit ferroptosis and restore progenitor cell reparative capacity represents a high-impact therapeutic approach.

## Methods

Sex was considered as a variable. Human samples included both males and females. Mouse smoke exposure models utilized male and female mice.

### Human lung tissue and plasma samples

Human lung tissue samples were obtained from the Thoracic Tissue Core at the University of Pittsburgh (supported by P30 DK072506, NIDDK, and the CFF RDP to the University of Pittsburgh). Donor lung samples were obtained from the Center for Organ Recovery and Education (CORE) at the University of Pittsburgh under Institutional Review Board (IRB) protocol CORID #300/#451. Donor lung samples originated from lungs deemed unsuitable for organ transplantation. All COPD samples were from lungs explanted from COPD patients that had undergone lung transplantation through the Thoracic Tissue Core (IRB #19100326). Lung tissues were stored at -80 °C until use or processed for paraffin embedding. All samples were deidentified prior to acquisition for experimental testing.

### Animal Studies and the Cigarette Smoke Exposure Model

All animal studies were approved by the Institutional Animal Care and Use Committee of the University of Pittsburgh. Mice were housed at the University of Pittsburgh and given *ad libitum* access to food and water as per standard protocols. Ant2 Cre-lox mice (Ant2^Fl/Fl or Fl/Y,^ Cre recombinase excision of exons 3 and 4 flanked by *loxP* sites) were a generous gift from Dr. Douglas Wallace (78). Ant2 flox mice were crossed with either constitutively active Sftpc*-*Cre–transgenic mice (SftpcCre^(Blh)^, generous gift from Dr. Jonathon Alder via Dr. Brigid Hogan (79) or AT2-cell specific SftpcCre^ERT2^ (stock no. 028054, Jackson Laboratories) and *Rosa26^membrane-targeted TdTomato/membrane-targeted GFP^*(*Rosa26*^mT/mG^) reporter mice (stock no. 007676, Jackson Laboratories). Ant2^WT,^ *Rosa26*^mT/mG^, Sftpc-Cre^+^ or Sftpc-CreERT2^+^ mice were used as control mice. All mice were maintained on a C57BL/6J background with backcrossing 10 generations. Genotyping of mice was performed using PCR primers specific for the Ant2 flox allele, Cre, Cre-ERT2 and Rosa26^mTmG^ transgenes for all mice. For the *Ant2* flox allele, the *Ant2* locus was genotyped using the forward primer 5’ ACTCAACCTAGGGCCTTGTG 3’ and the reverse primer 5’ GGGAGCATTCCTGAAAAATAA 3’ (35 cycles of PCR: 94° C for 20 secs, 56° C for 30 secs, and 72° C for 40 secs) to detect the *loxP* insertion. The targeted *Ant2* locus generated a 485 bp product while the wildtype locus gave a 354 bp product. Sftpc-CreERT2^+^ were treated with five days of 1mg/ml tamoxifen intraperitoneal injections.

To generate the ROSA26-Tet-On-Slc25a5 mouse model, a conditional and inducible knock-in targeting vector was designed. The backbone vector was constructed using the pROSA26-PA and pBigT plasmids (gifts from Frank Costantini; Addgene plasmids #21271 and #21270) (80). The targeting vector contains the following elements in 5’ to 3’ order between the two ROSA26 arm of homology: a splice acceptor, a loxP-flanked transcriptional STOP cassette (three tandem SV40 polyA terminators), the tetracycline-controlled transactivator rtTA/Tet-On 3G (81) with a bovine growth hormone polyA terminator, the tetracycline-responsive promoter TRE3G (82), and the C-terminally HA-tagged Slc25a5 (NM_007451.4) coding sequence linked in-frame via a T2A self-cleaving peptide to the fluorescent reporter mNeonGreen (83). In this configuration, Tet-On 3G expression from the ROSA26 promoter is blocked by the upstream Lox-Stop-lox cassette until Cre recombinase-mediated excision occurs, at which point doxycycline administration drives Slc25a5 expression in a tissue-specific and dose-dependent manner. The new strain was design, produced and genotyped by the Innovative Technologies Development Core and the Mouse Embryo Services Core in the Department of Immunology at University of Pittsburgh. For gene targeting, C57BL/6J zygotes were injected by pronuclear microinjection with a mixture containing EnGen Cas9 protein (0.67 µM; New England Biolabs), an sgRNA targeting the ROSA26 locus sgRosa26-1 (42 ng/µl ≈ 1.33 µM) (84) and the targeting vector (10 ng/µl). The single guide RNA was synthesized by in vitro transcription from a PCR generated template as previously described (85). Injected zygotes that developed to the 2-cell stage were transferred to pseudopregnant CD1 surrogate females. Potential founder pups were genotyped by PCR using a primer pair flanking the 5’ homology arm junction as well as two additional primer pairs spanning the Tet-On 3G, TRE3G, Slc25a5, and mNeonGreen sequences. The sequence of the PCR products from founders were verified by Sanger sequencing. Two founders carrying the correctly integrated transgene were identified and used to establish independent lines on the C57BL/6J background.

Male and female mice at 10-12 weeks of age (n = 9-10 per group) were subjected to the smoke of 4 unfiltered cigarettes per day (lot# 1R6F; University of Kentucky, Lexington, KY), 5 days a week for a duration of 6 months, using a smoking apparatus that delivers targeted cigarette smoke to single mice isolated in individual chambers as previously described (39,46). The control animals in each group were exposed to room-air alone. These mice were caged separately but housed in the same facility as their smoke-exposed counterparts. For lung harvesting, mice are euthanized with intraperitoneal pentobarbital overdose (200mg/kg). For pulmonary function testing, mice are fully anesthetized with ketamine/xylazine (100 and 15 mg/kg). The trachea is cannulated, and mice are mechanically ventilated. After anesthesia, mice receive pancuronium as a paralytic before pulmonary function tests. Mice underwent pulmonary function testing with SCIREQ flexiVent for in vivo respiratory mechanics. Bronchoalveolar lavage (BAL) was completed with 2 aliquots of 0.8mL PBS installation and removal with downstream processing for cell cytospins and BAL supernatant collection. For morphology assessment, lung lobes were inflated with 10% buffered formalin at a constant pressure of 25 cm H_2_O for 10 minutes. The lungs were then ligated, excised, and fixed in formalin for 18-24 hours before washing in PBS, storing in 70% ethanol and embedding in paraffin. Serial midsagittal sections were obtained for histological analysis at 5 µm thick.

### Mouse precision-cut lung slices (PCLS)

Mice were euthanized, and the lungs were perfused with ice-cold PBS through the right ventricle and pulmonary arteries. Lungs were inflated via the trachea with low–melting-point agarose (2% in PBS) and allowed to solidify on ice. The agarose-inflated lungs were sectioned into 300-µm slices using a vibratome. Slices were transferred to cell culture plates and maintained in DMEM/F-12 culture medium supplemented with 0.1% FBS and Penicillin-Streptomycin at 37°C with 5% CO2. PCLS were rested for 24 hours prior to experimental treatments. PCLS were exposed to CS for 24 hours.

### Mouse lung morphology analysis

Mouse lungs that were inflation-fixed as described were imaged with brightfield at 20x magnification. Images were masked to block out airways, blood vessels, and intra-alveolar immune cells or debris. Using an ImageJ based script we called “WaffleFry”, each masked image was overlaid with horizontal and vertical lines interspaced at 10-pixel intervals. Chord length measurements were automatically processed vertically and horizontally across this image grid. A chord length measurement was recorded when the subsequent pixel had tissue density (rather than airspace). Chord lengths of under 8 pixels were discarded to prevent capture of non-alveolar spaces. This analysis tool is similar to previously validated programs (39,46). It has been uploaded to Github at https://github.com/ckliment/WaffleFry.git.

### Human lung single cell RNA sequencing and analysis

Lung samples from COPD patients (GOLD stage IV, n=6, ages 58-68, 5 males and 1 female) and normal non-smoker donor controls (n=4, ages 56-68, 3 males, 1 female) were enzymatically digested as previously described (86) enriched EpCAM+ lung cells of the distal lung were separated out, and single-cell RNA sequencing was performed on the EpCAM+ lung cells. The sequencing results were analyzed using the Cell Ranger pipeline from 10x Genomics (v3.1.0, STAR v2.5.3a) and the Scanpy package (v1.8.0) (87). Reads were aligned to a hg38 human reference genome (GRCH38.97). Barcodes with fewer than 400 or more than 20,000 detected transcripts were excluded from the single-cell RNA library. Cells with a high proportion of mitochondrial-encoded transcripts were excluded. Cells with high background mRNA contamination were detected using the R library package SoupX (87) and also excluded from analysis. Variable genes were selected and ranked, and 3,426 genes identified as occurring in at least three samples were used as input for principal component analysis. After pre-processing, cells were clustered using standard cell markers. AT2 cells were specifically identified using cells positive for surfactant proteins, SFTPC and SFTPB, rather than Epcam to improve cell identification. Differential gene expression was calculated following leiden cell clustering and batch alignment with BBKNN (88). UMAPs were generated in Seurat (88). UMAPs and dotplots showing gene expression were generated via scanpy’s pl.umap() and pl.dotplot() function, respectively.

### Tissue and cell immunofluorescent staining

Immunofluorescent staining of lung tissue and cells in culture was completed as previously described (39,89). Lung tissue from human (normal non-smoker control, smoker without COPD, and COPD lungs) and mouse lung (air and CS exposed) were formalin fixed, paraffin embedded, and sectioned at 5 µm. Prior to staining and analysis, tissue sections were deparaffinized and rehydrated by a series of xylene and ethanol washes. Antigen retrieval was completed using sodium citrate buffer, pH 6.0, at 95°C. Tissue was permeabilized with 0.3% Triton X-100 and 1% BSA in PBS and blocked with 2% BSA in PBS. Human and mouse lung sections were stained for: ANT1 (Sigma-Aldrich #SAB2108761 rabbit polyclonal, 1:100), ANT2 (Abcam #192410 rabbit polyclonal, 1:100), rabbit anti-SPC (Sigma-Aldrich, #AB3786, 1:500), HTII-280 (Terrace Biotech, #TB-27AHT2-280), rat anti-LAMP3 (Novus Biologicals, Cat# DDX0191P, 1:200), podoplanin (Developmental Studies Hybridoma Bank, Univ. of Iowa, Anti-Pdpn #8.1.1), chicken anti-GFP (Aves Labs, #GFP-1010, 1:3500), rabbit anti-8-OHDG (Bioss, #BS-1278R, 1:200), rabbit anti-nitrotyrosine (Invitrogen, #A21285, 1:200), rabbit anti-HNE (MilliporeSigma #393207, 1:200), anti-Mitoferrin-1 (Invitrogen #PA5-119913, 1:250), chicken anti-HSP60 (Invitrogen #PA5-143571, 1:250), anti-mNeonGreen (ThermoFisher #NT-250, 1:1000) and secondary Alexa fluorophore antibodies (Molecular Probes). All tissue sections were stained with DAPI at 10 μg/mL for 10 minutes. Sections were mounted with Prolong Gold (Molecular Probes, ThermoFisher) and cured for at least 24 hr at 4°C prior to imaging. Images were obtained at the Center for Biologic Imaging (CBI) at the University of Pittsburgh on a Nikon A1R confocal microscope using 20 or 60x objective. Cell counts and image fluorescent intensities was quantified from 10-12 independent images per mouse using ImageJ Software (National Institutes of Health Open Access Software) and Nikon NIS elements.

### Transmission Electron Microscopy (TEM)

TEM imaging was completed on primary AT2 cells isolated from wildtype and ANT2-null mice (AT2-cell specific loss) as described. Cell processing for TEM was completed through the University of Pittsburgh Center for Biological Imaging using standard protocols. Analyzed mitochondrial size and area using Image J (90).

### Primary mouse alveolar type 2 cell isolation and 3D alveolar organoid cultures

Primary alveolar type 2 epithelial cells (AT2) were isolated from mouse lung as previously published (91,92) using negative (CD31 and CD45) and positive (EpCAM) bead selection (Miltenyi Biotech Micro Beads). Briefly, mice were euthanized, and lungs were perfused with ice-cold PBS, inflated, and enzymatically digested with dispase (Corning® #354235, 50 U/ml) followed by mechanical dissociation. Single-cell suspensions were filtered through strainers, subjected to red blood cell lysis followed by negative and positive bead selection using MAC columns, according to the manufacturer’s instructions. Cells were isolated by FACS sorting for GFP+ cells. Isolated AT2 cells were used directly for downstream assays, including RNA isolation, plating for Seahorse analysis, and alveolar organoid seeding.

For 3D alveolar organoids, isolated AT2 cells were plated in Matrigel (BD Biosciences, #354230) at 3,000 cells per dome in glass-bottom 96-well plates (Cellvis, Cat#: P96-1.5H-N) according to published protocols using a fibroblast-free system. Organoid growth was monitored over 14 days, with media changes every other day. Organoids were stained with calcein for 30 minutes according to the manufacturer’s protocols immediately prior to imaging. Organoids were imaged using the BioTek Cytation Live Cell Imager at 10x magnification or using Crest Deep SIM at 4x magnification. Organoid number and area were quantified on Day 14. Organoids on 96-well plates were washed with 1x PBS, fixed with 4% PFA, and stored in 5% BSA in PBS at 4°C.

### Human cell culture

The human bronchial epithelial cell line (BEAS-2B) were obtained from ATCC. Cells were cultured on tissue culture-treated polystyrene plates and were propagated when 70-90% confluent. BEAS-2B cells were maintained in Dulbecco’s modified Eagle media (DMEM) with Nutrient Mixture F-12 and supplemented with 5% fetal bovine serum and 1% penicillin/streptomycin. Cells were propagated by trypsinization with 0.25% trypsin-EDTA (Gibco #25200-056) and neutralized by trypsin neutralizing solution (Gibco #R-002-100).

### Targeted gene suppression and CRISPR-Cas9 gene knockout

BEAS-2B cells were grown to 25-50% confluency and then transfected with 150 nM siRNA ON-TARGETplus smart pools (Dharmacon: ANT2 (*SLC25A5*, #L-007486-02-0005) and non-targeting control pool (#D-001810-10-05) by lipofection with Lipofectamine 3000 (Invitrogen) using the manufacturer’s protocol and as previously described (89). Briefly, lipid-siRNA complexes were formed in Opti-MEM Reduced Serum Medium (ThermoFisher) at room temperature for 20 minutes, then slowly dripped onto the cells. ANT2 was knocked down for 48-72 hours prior to any additional treatments or cell collection.

BEAS-2B cells were used to create Crispr knockout clones for *SLC25A5* (ANT2). All lentiviruses were packaged in HEK293 cells by co-transfection of the lentiviral vector and packaging plasmids (pCMV-delta8.9 (Expressing HIV gag/Pd, Rev, and tat) and pCMV-VSV.G (to express envelope protein)) as previously described (93) in HEK293 cells. Crispr clones were created as previously described (89). The LentiCRISPR v2 plasmid (AddGene #52961) was used to create viral vectors expressing specific sgRNA guides and Cas9. The sgRNA guides designed included: SLC25A5 sgRNA guide 1: 5’-TGGCATCGGGTGGTGCCGCA*GGG* -3’; SLC25A5 sgRNA guide 2: 5’-GGCATCGGGTGGTGCCGCAGGGG-3’; and scrambled sgRNA guide: 5’- GCGTGGCGTACCGCATACCA-3’. Following viral transduction, individual cell clones were selected in the presence of 1.5 ug/ml puromycin by plating single cells per well in a 96-well plate. Clones were expanded and individually tested for ANT2 knockout by Western blot analysis.

### Reverse Transcription-quantitative PCR

Mouse lung tissue and culture cells were homogenized in Trizol, total RNA was isolated according to the manufacturer’s instructions (Thermo Fisher, Grand Island, NY), and RNA was reverse transcribed (Applied Biosystems, Grand Island, NY). Real-time PCR was performed using total cDNA and primer/probe pairs flanking introns specifically targeted the genes of interest using the SSO Advanced (Bio-Rad) master mix for real-time PCR on a Bio-Rad CFX96 Real-Time PCR machine. The PCR protocol included cDNA synthesis at 50°C for 15 min, followed by repeat cycles of inactivation at 95°C for 15 min, denaturing of DNA at 95°C for 15 seconds, annealing at 60°C for 30 seconds, extension at 72°C for 30 seconds (repeat for a total of 40 cycles). Targeted TaqMan FAM probe sets included mouse *Gapdh* (Mm99999915_m1), *Slc25a4* (Mm01207393_m1), *Slc25a5* (Mm00846873_g1) from ThermoFisher. The real-time PCR data were analyzed by the ΔΔCT method with normalization to *Gapdh* and controls.

### Metabolic Assessment of primary murine AT2 cells and in vitro cell cultures

The Seahorse MitoStress assay (XF96 Flux Analyzer, Agilent) was used to perform metabolic analysis of BEAS-2B cells with ANT2 modulation and of isolated primary murine AT2 cells. Cells were cultured at a density of 80,000-100,000 cells per well in 96-well Seahorse assay plates (Agilent) as stable CRISPR clones or isolated AT2 cells. Prior to the assay, cells were washed, followed by washes in buffered Seahorse Assay medium (pH 7.4), and subsequent metabolic testing was performed according to the manufacturer’s protocol (Agilent Seahorse XF96). Oligomycin (2 μM, ATPase inhibitor), FCCP (0.25 μM mitochondrial uncoupler), and a cocktail of rotenone (0.5 μM, ETC complex I inhibitor) and antimycin A (0.5 μM, ETC complex III inhibitor) were sequentially injected after three basal rates were measured. Real-time oxygen consumption rate (OCR) is determined, including basal OCR (before test kit inhibitors), maximal OCR (after FCCP treatment, a mitochondrial uncoupler), ATP production (OCR after injection of the complex V inhibitor oligomycin), and proton leak. Sample measurements were normalized to cell density, as determined by the CyQUANT assay according to the manufacturer’s protocol. The XF ATP Rate Assay (XF96 Flux Analyzer, Agilent) was used per the manufacturer’s instructions as follows. Cells were cultured at a density of 80,000-100,000 cells per well in 96-well Seahorse assay plates (Agilent) as stable CRISPR clones Data were analyzed in the Agilent ATP Assay Report Generator, and statistics were analyzed in Prism 10.1.0. NAD^+^/NADH and steady-state ATP assays (luciferin-luciferase bioluminescence ATP Determination Kit) were performed according to the manufacturer’s protocols (ThermoFisher Scientific) using 100,000 cells per sample.

### Flow Cytometric Analysis of Mitochondrial Function

Mitochondrial membrane potential in cells was determined using the TMRM (Invitrogen, Catalog No. I34361), per the manufacturer’s protocol. Mitochondrial superoxide generation in cells was determined using the MitoSox Red mitochondrial superoxide indicator (Invitrogen, Catalog number M36008), per the manufacturer’s protocol. Extracellular Hydrogen Peroxide generation in cells was determined using the Amplex Red indicator (Invitrogen), per the manufacturer’s protocol.

### GSS/GSSG Analysis

GSH generation in cells was determined using the GSH/GSSG assay (Abcam, ab138881) per the manufacturer’s protocol. Cell lysates were incubated with Thiol Green indicator to quantify reduced glutathione (GSH), while parallel samples were incubated with Thiol Green and GSSG Probe to measure total glutathione (GSH + GSSG). Fluorescence was measured at Ex/Em = 490/520 nm, and concentrations were calculated from glutathione standard curves.

### Iron Quantification

The intracellular content of Fe^2+^ was measured using the Iron Assay Kit (cat. no. ab83366; Abcam). The cells were collected and homogenized with iron assay buffer on ice. The samples were then centrifuged at 13,000 × g for 10 min at 4°C, and 300 µl supernatant was collected. Iron reducer (300 µl) was added, mixed, and incubated at room temperature for 30 min. A volume of 200 µl of iron probes was then added and mixed thoroughly. The reaction mixture was incubated for 30 min at room temperature away from light. The absorbance was measured on a colorimetric microplate reader at 593 nm. Intracellular and mitochondrial iron were analyzed using FerroOrange and MitoFerroGreen dyes (Dojindo, F374 and M489), respectively, according to the manufacturer’s protocols. Cells were incubated with 1 μmol/L dye solution and incubated for 30 minutes. Viable cells were then analyzed by flow cytometry.

### Heme Assay

Cells were plated at 0.5×10^6 cells/mL in a 10 cm culture dish. After removing media and washing with PBS, cells were scraped and pelleted by centrifugation at 5000×g for 5 minutes at 4°C. BAL samples were centrifuged at 1000×g for 5 minutes. The pellet was washed, re-pelleted, transferred to an iron-free 1.5 mL Eppendorf tube, and divided into two equal aliquots. Lung tissue was lysed. For each aliquot, 500 µL of 2M oxalic acid was added. A portion was used for protein determination by BCA analysis. One aliquot was kept at room temperature (background), and the other was heated at 95°C for 30 minutes to release iron from heme and produce PPIX. Post-heating, samples were centrifuged at 1000×g for 10 minutes at 4°C. Supernatants were transferred into a black 96-well plate. Fluorescence was measured at excitation 400 nm/emission 620 nm. The fluorescence from unheated samples was subtracted from heated samples. Using the relation 1 µM heme = 15,200 fluorescence units at 622 nm, heme concentrations were calculated and normalized to protein content (94). Heme concentrations were calculated using the same fluorescence-to-heme conversion factor.

### Bulk RNA sequencing and real-time PCR

RNA was isolated from primary mouse AT2 cells and the Beas2b cell line and submitted to Novogene Inc. for transcriptomic analysis and quantitative real-time PCR (qRT-PCR) validation. Bulk RNA sequencing was performed by Novogene Inc. using an Illumina sequencing platform based on sequencing-by-synthesis (SBS) technology. Raw sequencing reads were subjected to quality control and filtering before alignment to the mouse reference genome (mm10) for murine samples or the human reference genome (hg38) for BEAS-2B cell samples.

### Western blot analysis

Tissue and cell lysates were prepared using RIPA buffer supplemented with DNAse I (ThermoFisher) and a protease inhibitor cocktail (Cell Signaling). Protein concentration was determined by the Bradford assay (Pierce). Proteins were separated by 4-20% gradient polyacrylamide gel electrophoresis (Bio-Rad) and transferred to a nitrocellulose membrane using a semi-dry blot transfer system (Bio-Rad). Blots were blocked with 5% bovine serum albumin (ThermoFisher) for 1 hour, then immunoblotted with the primary antibody overnight at 4°C. Primary antibodies used were targeted against ANT1 (Sigma-Aldrich #SAB2108761), ANT2 (Abcam #192410), NCOA4 (Abcam #Ab86727), FTH (Abcam #Ab65080), SLC7A11 (Abcam #Ab37185), GPX4 (Abcam #Ab125066), TFRC (Cell Signaling #46222T), FPN (Invitrogen #PA5-22993), MFRN1 (Invitrogen #PA5-119913), MFRN2 (Bioss #BS-7157R), HMOX1 (Cell Signaling #43966). Secondary antibodies used were Invitrogen GOXMO HRP high XADS (#A16078, 1:10,000). Blots were developed using SuperSignal West Pico PLUS Chemiluminescent Substrate (Thermo Scientific #34580) and imaged with a ChemiDoc XRS+ (Bio-Rad) detector.

### Assessment of lipid peroxidation using C11-BODIPY (581/591)

A total of 150,000 cells per well were seeded on 12-well plates. The next day, cells were treated with or without CSE and with ferroptosis inhibitors, liproxstatin (250 nM, Selleckchem, S7699) or Deferoxamine (50 μM, Sigma-Aldrich D9533), for 24 hours. The following day, cells were treated with (1 S,3 R)-RSL3 (250 nM, Cayman Chemical N0.19288) to induce ferroptosis for 6 h. Cells were incubated with C11-BODIPY (581/591) (1 μM, Invitrogen D3861) for 20 min at 37 °C, then washed with PBS and harvested by trypsinization. Subsequently, cells were resuspended in 100 μL of fresh PBS and analyzed using the 488-nm laser of a flow cytometer (LSR Fortessa, BD Biosciences). Cell death is measured using Zombie NIR (Biolegend, Cat# 423105), according to the manufacturer’s instructions. At least 10,000 events were analyzed per sample, and data were collected from the FL1 detector (C11-BODIPY) with a 488nm laser, 505LP and 530/30 BP filter, and (ZOMBIE NIR) with 647 nm laser, 750LP and 780/60 BP filter. Data were analyzed using FlowJo Software (v8.4.1).

### Statistical Analysis

Mean densitometry (Image Lab Software) and all other quantitative data (mean ± SEM) were normalized to appropriate control groups. Statistical analyses were completed using GraphPad Prism 10.1.0. Data were assessed for sample distribution. If samples were normally distributed, then the data were analyzed using ANOVA with Fisher’s LSD post-test. If the data were not normally distributed, nonparametric analyses, including Kruskal-Wallis, Tukey’s, and/or Mann-Whitney, were used. For all statistics, a p-value less than 0.05 was considered to be statistically significant. The number of replicates for each experiment is specified in the figure legends.

## Acknowledgments

A special thank you to Dr. Steven Shapiro for his guidance and input. We thank Maggie Sedgwick, Andy Metz, Sahana Krishna Kumaran, Adriana Leme, and the Center for Biological Imaging (CBI) for their technical support. We thank the funding agencies that have supported this work.

## Data and materials availability

All data and code associated with this study are available in the main text or the supplemental materials. Contact Corrine Kliment for study correspondence and material requests.

## Author contributions

Conceptualization: CRK, UM

Computational analysis, experimental assays and data processing: UM, JS, SG, NT, HX, CRK, YZ, QH, MK

Funding acquisition: CRK, UM, MK

Writing – original draft: CRK, UM

Writing – review & editing: CRK, UM, MK

## Funding

National Institutes of Health grant – NHLBI: K08HL141595 (CRK), NHLBI: R01HL168050-01 - (CRK), NHLBI Diversity Supplement to R01HL168050-01 (UM), NIA R61AG095927 (CRK, MK), R01 HL141380 (MK)

Burroughs Wellcome Fund Career Award for Medical Scientists (CRK)

Parker B Francis Pulmonary Fellowship (CRK)

UPMC Immune Transplant and Therapy Center, ITTC grant (CRK)

LongFonds BREATH Consortium (MK)

