## Supplemental Figures S1-6 for "Adenine nucleotide translocase 2 (ANT2) deficiency reprograms ferroptosis in alveolar progenitor cells to promote emphysema"

Mbaekwe et al.

Supplemental Figure 1:

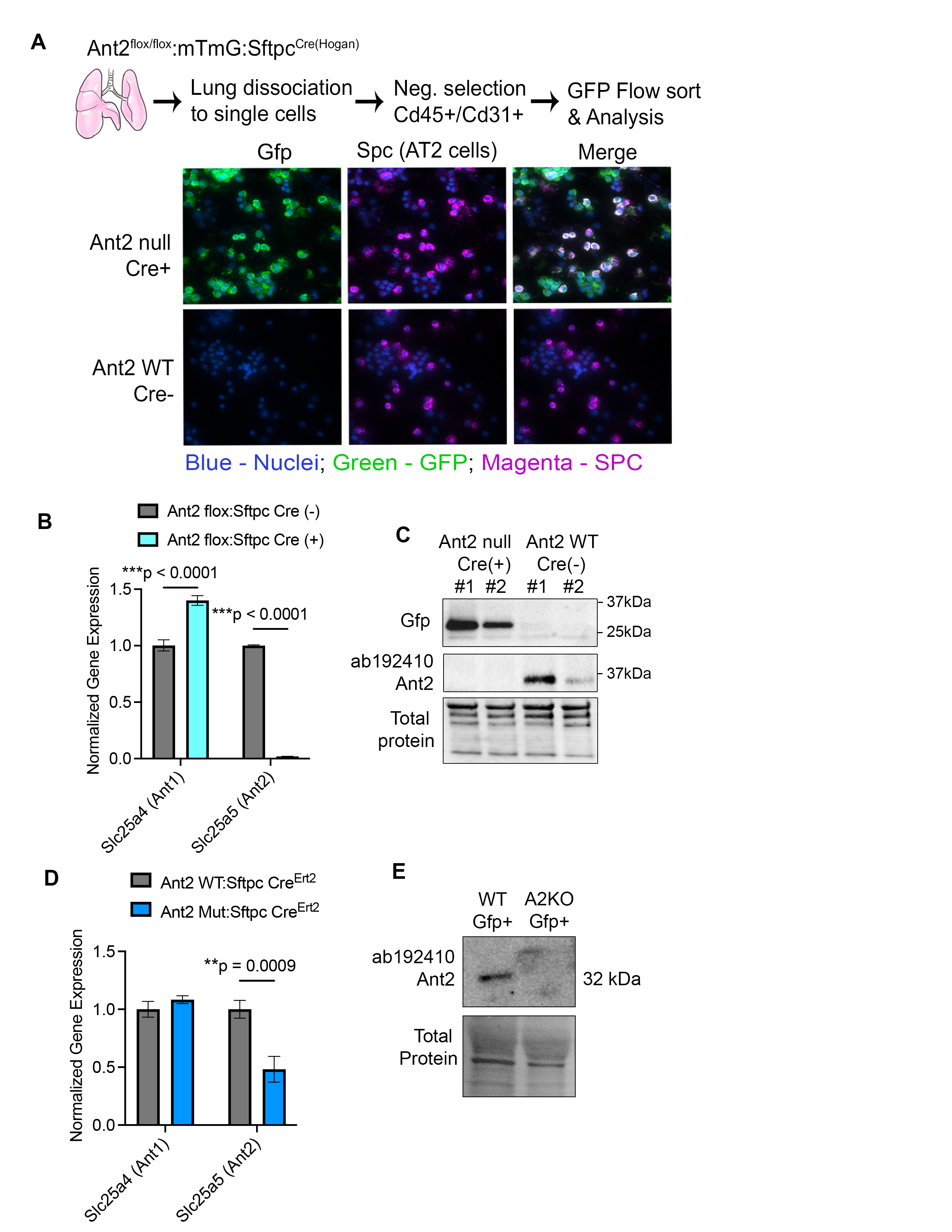

**Figure S1: Confirmation of Ant2 KO in SPC and ERT2 mice.**

(A) Schematic of AT2 cell isolation. Lungs are dissociated into single-cell suspensions and subjected to a CD45/CD31-negative selection with GFP flow sorting. According to the immunofluorescence of isolated AT2 cells. Nuclei are indicated in blue, GFP in green, and pro-SPC in magenta (B) Normalized Gene expression with *Slc25a4* and Slc25a5 from isolated AT2 cells from WT and Ant2-null^EPI^ mice. (C) Western blot of Ant2 protein and total protein blot from isolated AT2 cells from WT and Ant2-null^EPI^ mice. (D) Normalized Gene expression of *Slc25a4* and Slc25a5 from isolated AT2 cells from WT and Ant2-null^AT2^ mice (E) Western blot of Ant2 protein and total protein blot from isolated AT2 cells from WT and Ant2-null^AT2^ mice.

Supplemental Figure 2:

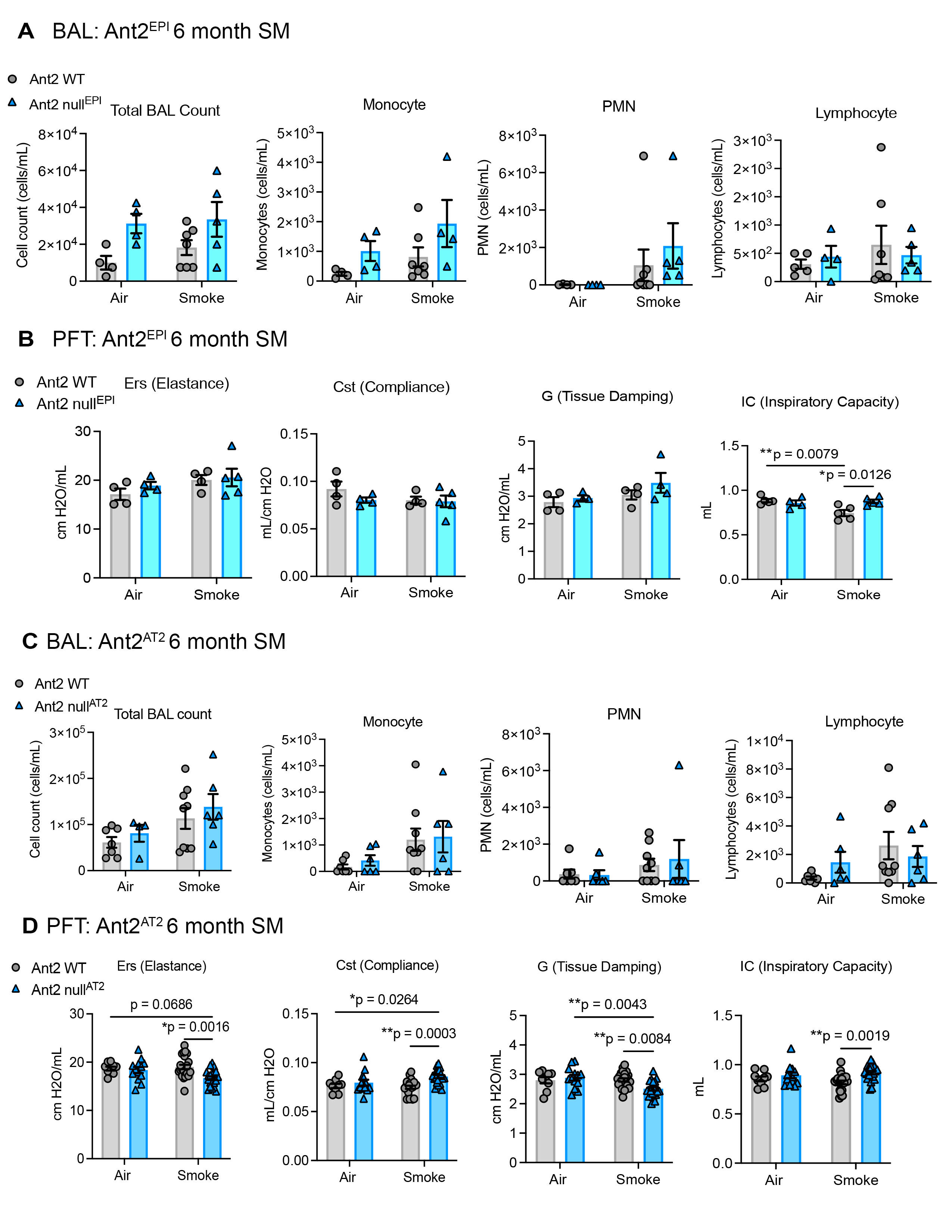

**Figure S2: Characterization of bronchoalveolar lavage and pulmonary function in ANT2 mouse models**

Lung mechanics were measured through oscillometry by the flexiVent (SCRIREQ). Respiratory system resistance (Rrs), respiratory system elastance (Ers), static compliance (Cst), inspiratory capacity (IC), airway resistance (Rn), tissue resistance/damping (G), tissue elastance (H) are shown (A) wild-type or Ant2-null ^ERT2^ mice treated with air or smoke (SM) for 2 months. (n = 6-8 mice per group) and (B) wild-type or Ant2-null ^ERT2^ mice treated with air or smoke (SM) for 6 months. (n = 9-10 mice per group). Data are represented as mean ± SEM. *p < 0.05, **p < 0.01, ***p < 0.0001, with Statistics by two-way ANOVA with Tukey’s post hoc test. P values are noted.

Supplemental Figure 3:

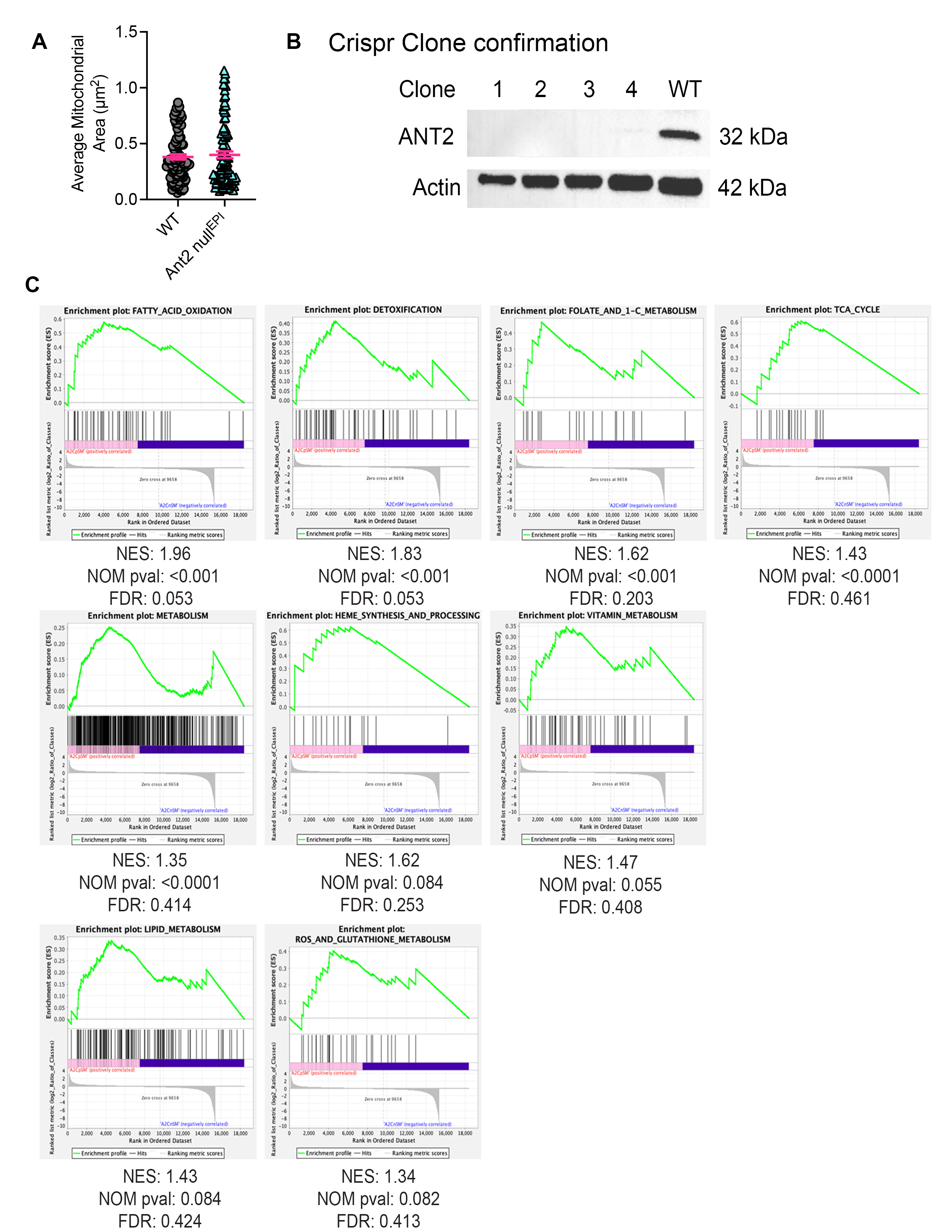

**Figure S3: Confirmation of ANT2 KO cells and bulk RNA sequencing GSEA analysis**

(A) Total BAL and differential BAL fluid cell counts (monocyte, polymorphonuclear leukocytes (PMN), lymphocytes) from wild-type (WT) or Ant2-null ^EPI^ mice treated with air or smoke (SM) for 6 months. (B) Total BAL and differential BAL fluid cell counts (monocyte, polymorphonuclear leukocytes (PMN), lymphocytes, macrophages) from wild-type (WT) or Ant2-null ^AT2^ mice treated with air or smoke (SM) for 2 months. (C) Average alveolar chord length quantification from histological images of wild-type (WT) or Ant2-null ^AT2^ mice treated with air or smoke (SM) for 2 months. Each point represents a biologically independent mouse tissue sample, with average alveolar chord lengths measured from 6-8 mice per group, and 12–14 images per mouse. (D) Total BAL and differential BAL fluid cell counts (monocyte, polymorphonuclear leukocytes (PMN), lymphocytes, macrophages) from wild-type (WT) or Ant2-null ^AT2^ mice treated with air or smoke (SM) for 6 months. Data are represented as mean ± SEM. *p < 0.05, **p < 0.01, ***p < 0.0001, with Statistics by two-way ANOVA with Tukey’s post hoc test. P values are noted.

Supplemental Figure 4:

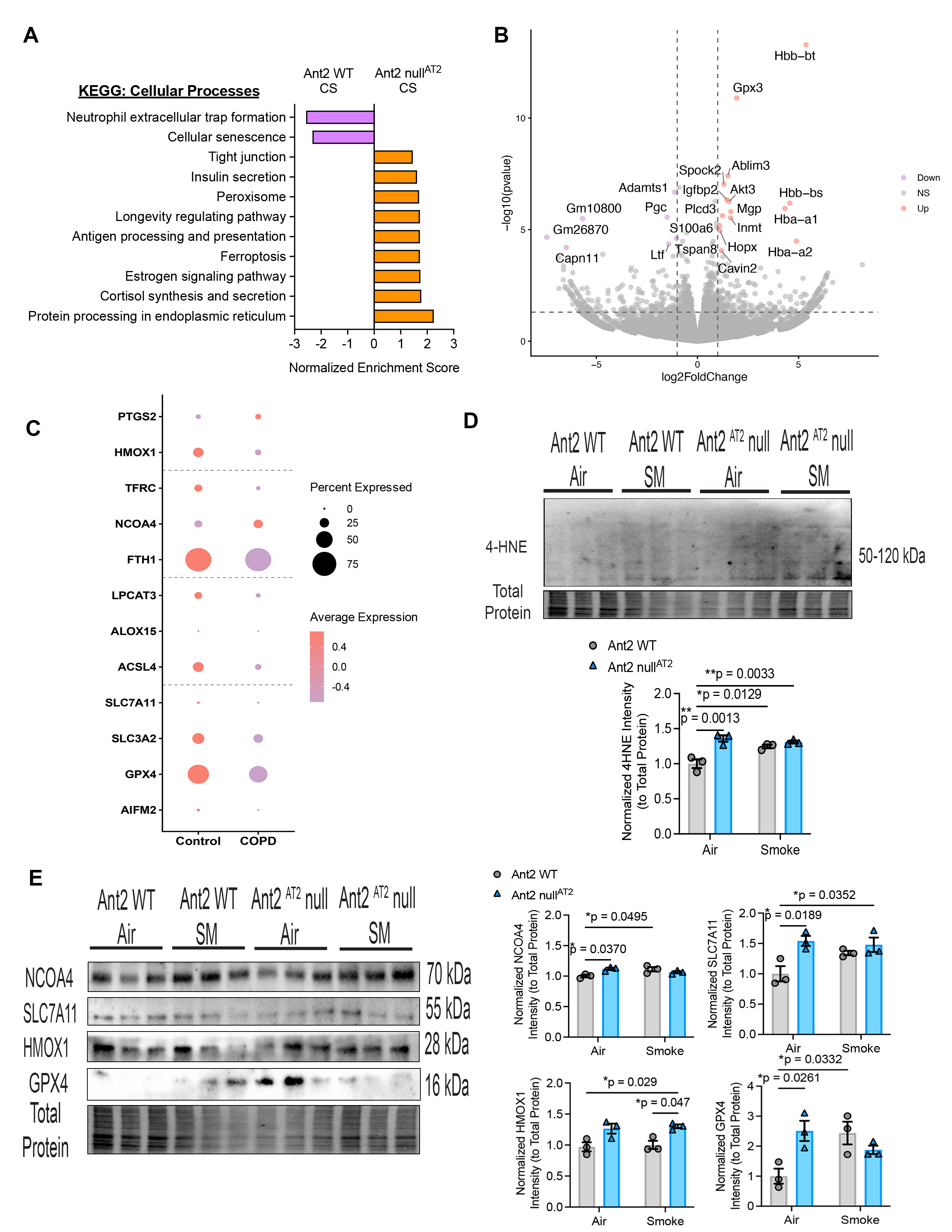

**Figure S4: Analysis of Ferroptosis-related genes in human and mouse lung**

(A) Gene set enrichment analysis (GSEA) pathway analysis matched to Kyoto Encyclopedia of Genes and Genomes enrichment category for Cellular Processes was performed on significantly differentially expressed gene lists (DEGs) generated comparing isolated AT2 cells from wild-type (WT) or Ant2-null ^EPI^ mice Beas2b cells treated with CS for 6 months (n =2-4 per treatment group). Significantly enriched pathways are indicated by (p-value < 0.05 and FDR < 0.05). (B) Volcano Plot of DEGs from isolated AT2 cells from wild-type (WT) or Ant2-null^AT2^ mice (n =3 per treatment group). Orange are up-regulated DEGs, purple are down-regulated DEGs, and grey are not significant DEGs. (C) Single-cell RNA sequencing results obtained on isolated lung epithelial cells from normal control (n=4) and COPD (n=6) subjects (86). Mean expression and fraction of cells with gene expression of ferroptosis genes for AT2 cell cluster. Circle size represents the percentage of cells expressing the gene, and color gradient represents the average expression in cells. (D) Western blot and respective quantification of total 4HNE from lung lysate from wild-type and Ant2-null^AT2^ mice treated in air or CS for 6 months. (E) Western blot for ferroptosis genes performed on cell lysates from murine WT and Ant2-null^AT2^ mice either treated in air or CS for 6 months. Respective quantifications. Data are represented as mean ± SEM. *p < 0.05, **p < 0.01, ***p < 0.0001, with Statistics by two-way ANOVA with Tukey’s post hoc test. P values are noted.

Supplemental Figure 5:

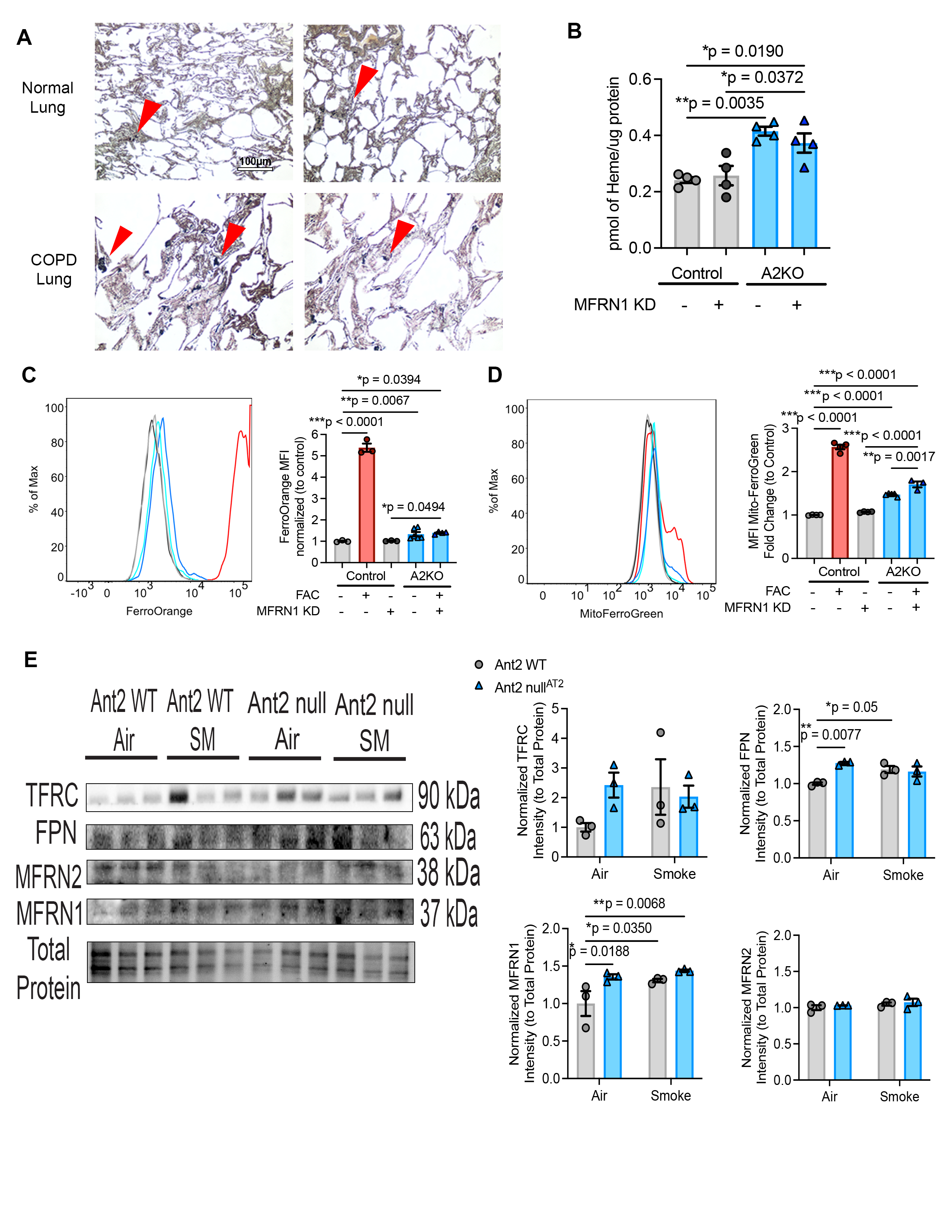

**Figure S5: Analysis of iron-related mechanisms through Mitoferrin-1 and in the murine smoking model.**

(A) Perls Stain for healthy and COPD tissue. Red arrows indicated iron deposits in lung tissue. (B) Heme content was determined by oxalic acid-induced conversion of heme to fluorescent protoporphyrin IX (PPIX) for CRISPR-Cas9 control and ANT2 KO Beas2b cells treated with or without siRNA knockdown of MFRN1. Background-corrected fluorescence (heated minus unheated samples) was normalized to total protein concentration. (C) Histogram of flow cytometry of control and ANT2 KO cells treated with or without siRNA knockdown of MFRN1 for FerroOrange (intracellular iron) and (D) MitoFerroGreen (mitochondrial iron) analyzed in flow cytometry. The bar graph shows the mean probe fluorescence intensity across at least 3 experiments (derived from 3 independent determinations). Ferric Ammonium Citrate (FAC)-treated groups served as a positive control. (E) Western blot of iron import and export genes in cell lysates from WT and Ant2-null^AT2^ mice treated with or without cigarette smoke for 6 months. Respective quantifications. Data are represented as mean ± SEM. *p < 0.05, **p < 0.01, ***p < 0.0001, with Statistics by two-way ANOVA with Tukey’s post hoc test. P values are noted.

Supplemental Figure 6:

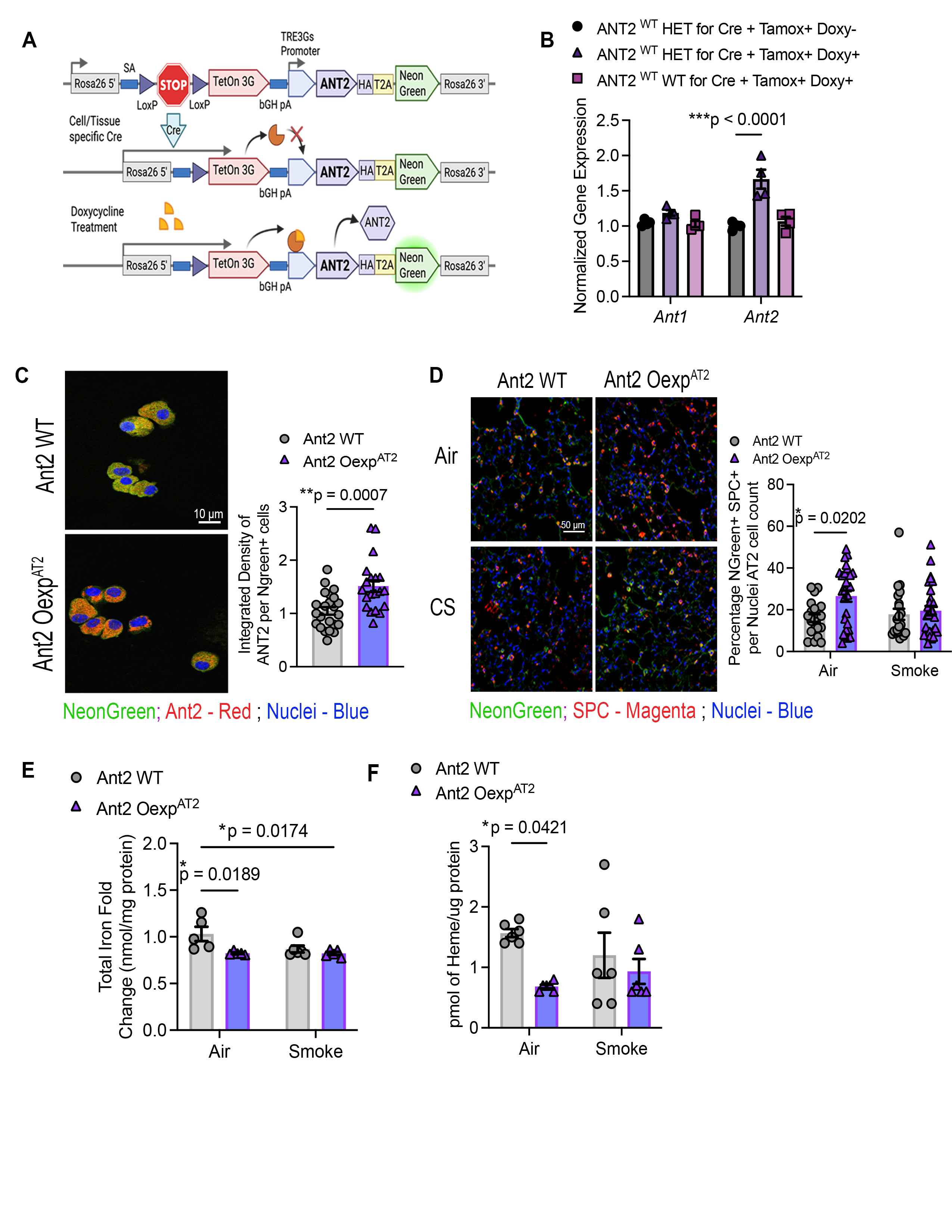

**Figure S6: Confirmation of overexpression of ANT2.**

(A) Schematic of Ant2^OExp_AT2^ mice with tissue-specific and temporal control of ANT2 expression using a Cre-recombinase and tetracycline-inducible system with treatment of doxycycline. (D) 3D alveolar organoids from wild-type or Ant2^OExp_AT2^ mice (B) Ant2^OExp_AT2^ mice TetOn RNA confirmation from whole mouse lung lysate. (C) Ant2^OExp_AT2^ mice TetOn protein and immunofluorescence confirmation from isolated AT2 cells from WT and Ant2^OExp_AT2^ mice. Immunofluorescence quantification of human SLC25A5 in isolated AT2 cells. (D) Immunofluorescence of wild-type or Ant2 Oexp ^AT2^ mice lungs treated with air or smoke (SM) for 6 months. (n = 3 mice per group). DAPI is indicated in blue, NeonGreen is indicated in green, and SPC is indicated in red. Respective quantification of NeonGreen+ cells across the entire lung. Scale bar is 50μm. (E) Iron assay quantification of WT and Ant2-Oexp^AT2^ mice treated with or without cigarette smoke for 6 months, detecting total iron. (F) Heme assay quantification in WT and Ant2^OExp_AT2^ mice treated with or without cigarette smoke for 6 months. Statistics are calculated using the unpaired nonparametric Mann-Whitney test and two-way ANOVA with Tukey’s post hoc test when appropriate. Data are represented as mean ± SEM. *p < 0.05, **p < 0.01, ***p < 0.0001.
